# UFMylation of Pyruvate Dehydrogenase Regulates Mitochondrial Metabolism

**DOI:** 10.64898/2026.03.18.712693

**Authors:** Phong T. Nguyen, Zheng Wu, Dohun Kim, Tamaratare Ogu, Siyu Yin, Varun Sondhi, Feng Cai, Trevor S. Tippetts, Annie Jen, Evgenia Shishkova, Ling Cai, Dennis Dumesnil, Margaret Cervantes, Hongli Chen, Prashant Mishra, Joshua J. Coon, Gerta Hoxhaj, Min Ni, Ralph J. DeBerardinis

## Abstract

The ubiquitin-fold modifier 1 (UFM1) post-translational modification (PTM), or UFMylation, regulates protein homeostasis and is essential for human development. Yet the roles of the de-UFMylase, UFM1-specific peptidase 2 (UFSP2), which removes UFM1 from UFMylated proteins, remain poorly characterized. Here, we demonstrate that UFMylation and UFSP2 regulate mitochondrial metabolism. Quantitative proteomics in UFSP2-deficient cells revealed the accumulation of many proteins previously unknown to be impacted by UFMylation. These included components of the mitochondrial ribosome, electron transport chain (ETC), and pyruvate dehydrogenase (PDH) complex. Functional analyses demonstrated that excessive UFMylation in UFSP2-deficient cells increases mitochondrial respiration, glucose oxidation in the tricarboxylic acid (TCA) cycle, and PDH enzymatic activity. We identified dihydrolipoamide S-acetyltransferase (DLAT), the E2 component of PDH, as a direct UFMylation substrate, with lysine 118 (K118) as the primary conjugation site. Mutating K118 to arginine (K118R) abolished DLAT UFMylation and reduced pyruvate oxidation, identifying this modification as an activator of PDH. These findings reveal a UFMylation-based regulatory mechanism that controls mitochondrial function by inducing utilization of pyruvate as a TCA cycle fuel.

## INTRODUCTION

The ubiquitin-like UFM1 post-translational modification (PTM), also known as UFMylation, is the attachment of the ubiquitin-like peptide UFM1 (Ubiquitin-Fold Modifier 1) to proteins using an E1 - E2 - E3 enzymatic cascade analogous to ubiquitination^1^. Unlike the hundreds of enzymes contributing to ubiquitination activity, UFMylation is much simpler, using single E1 (Ubiquitin-like modifier activating enzyme 5, UBA5), E2 (UFM1-conjugating enzyme 1, UFC1), and E3 (UFM1 ligase 1, UFL1) enzymes, plus two deUFMylases (UFM1-specific peptidase 1, UFSP1 and UFM1-specific peptidase 2, UFSP2)^2,3^. Dysregulated UFMylation is implicated in human diseases including neurodegeneration and cancer^4^. Mutations of genes involved in UFMylation cause developmental aberrations in humans^5–9^ and mice^10–18^.

Despite the well-characterized components of the UFMylation system, only a few targets have been identified, validated, and mechanistically studied^19^. Functional requirements for UFM1 conjugation have been demonstrated in ribosome quality control (RQC)^20–24^, selective degradation of the ER membrane (ER-phagy)^25^, DNA damage responses^26–28^, transcriptional regulation^29^, immune checkpoint regulation^30^, and ER stress protection^31,32^. In contrast, the biological function of UFSP2-mediated de-UFMylation, the process of removing UFM1 from UFMylated proteins, remains largely unknown.

UFSP1 and UFSP2 are cysteine proteases that share a conserved catalytical triad/tetrad consisting of cysteine, histidine, aspartate, and tyrosine residues^33–35^. Human UFSP1, previously mis-annotated as an inactive, short isoform, also exists as a longer isoform containing a functional catalytic domain^36–38^. UFSP1 is cytosolic and crucial for proteolytic cleavage of pro-UFM1 into the shorter, mature UFM1 that serves as the substrate for protein modification^36–38^. UFSP2 contains an N-terminal domain missing from UFSP1 and can localize to different organelles. For instance, interaction with the ER membrane protein ODR4 recruits UFSP2 to the ER^39^, where it mediates the deconjugation of ribosomal protein large subunit 26 (RPL26), a key UFMylated target within the large ribosomal subunit (60S) involved in RQC^24^. UFSP2 is also recruited to DNA double-strand breaks (DSB) following Ataxia-Telangiectasia Mutated (ATM) kinase-dependent phosphorylation^40^.

Several UFSP2 sequence variants are linked to monogenic disorders in humans. Pathogenic variants within the catalytic domain (e.g., Y280H, D426A, and H428R) are associated with Spondyloepimetaphyseal dysplasia, Di Rocco type (SEMDDR) and Beukes familial hip dysplasia and follow an autosomal dominant inheritance pattern^41–43^. In contrast, the V115E variant in the N-terminal domain causes an autosomal recessive developmental and epileptic encephalopathy^44^. Pathogenic variants in either the catalytic or N-terminal domain result in a loss of UFSP2 function and accumulation of UFMylated proteins.

In this work, we aimed to further define UFSP2’s regulatory functions by profiling the UFMylated proteome in UFSP2-deficient cells. Surprisingly, we observed an accumulation of many potentially UFMylated mitochondrial proteins in UFSP2-deficient cells and found that these cells display elevated mitochondrial respiration and activity of the pyruvate dehydrogenase complex (PDH). Mechanistic studies revealed that UFSP2 deficiency results in hyper-UFMylation of the PDH E2 subunit, dihydrolipoamide S-acetyltransferase (DLAT), enhancing the conversion of pyruvate to acetyl-CoA by PDH. These findings establish UFSP2 as a regulator of glucose oxidation within the mitochondria.

## RESULTS

### UFM1 immunoprecipitation yields many mitochondrial proteins in UFSP2-deficient cells

To identify candidate UFMylated proteins, we engineered UFSP2-knockout 293T cells that stably express either Flag-tagged wildtype UFM1 (UFM1^WT^) or a conjugation-deficient UFM1 mutant (UFM1^ΔGSC^). The latter lacks the terminal three residues (GSC) whose glycine is required for protein target conjugation (Fig. 1A). To maximize detection of UFMylated proteins, we transiently overexpressed the core UFMylation machinery: UBA5, UFC1, UFL1 and DDRGK1 (UFBP1) as previously described^45,46^. Immunoprecipitation (IP) using Flag-M2 affinity resin isolated UFMylated proteins and their associated complexes (Fig. 1B). Mass spectrometry (MS) revealed many UFMylated candidates in the UFM1^WT^ eluate compared to the UFM1^ΔGSC^ control (adjusted p-value <0.05, fold change > 2, Fig. 1C). Enriched UFMylated proteins in the UFM1^WT^ sample included enzyme components of the UFMylation pathway and known UFMylation targets, validating the approach (Fig. 1C). The analysis also revealed a large number of mitochondrial proteins among enriched candidates in cells expressing UFM1^WT^ (91 out of 639 proteins, Fig. 1C, Extended Fig. 1). These included mitochondrial ribosomal proteins (Extended Fig. 1A), components of electron transport chain (ETC) complexes I-V (Extended Fig. 1B-F), and the PDH complex (Fig. 1C). UFMylated proteins have been found in the ER^20–24^, nucleus^47^, and cytosol^30,48–50^, but not the mitochondria. The large number of mitochondrial proteins enriched in UFM1^WT^ IP-MS raised the possibility that UFMylation has regulatory effects over mitochondrial function.

**Fig. 1:**
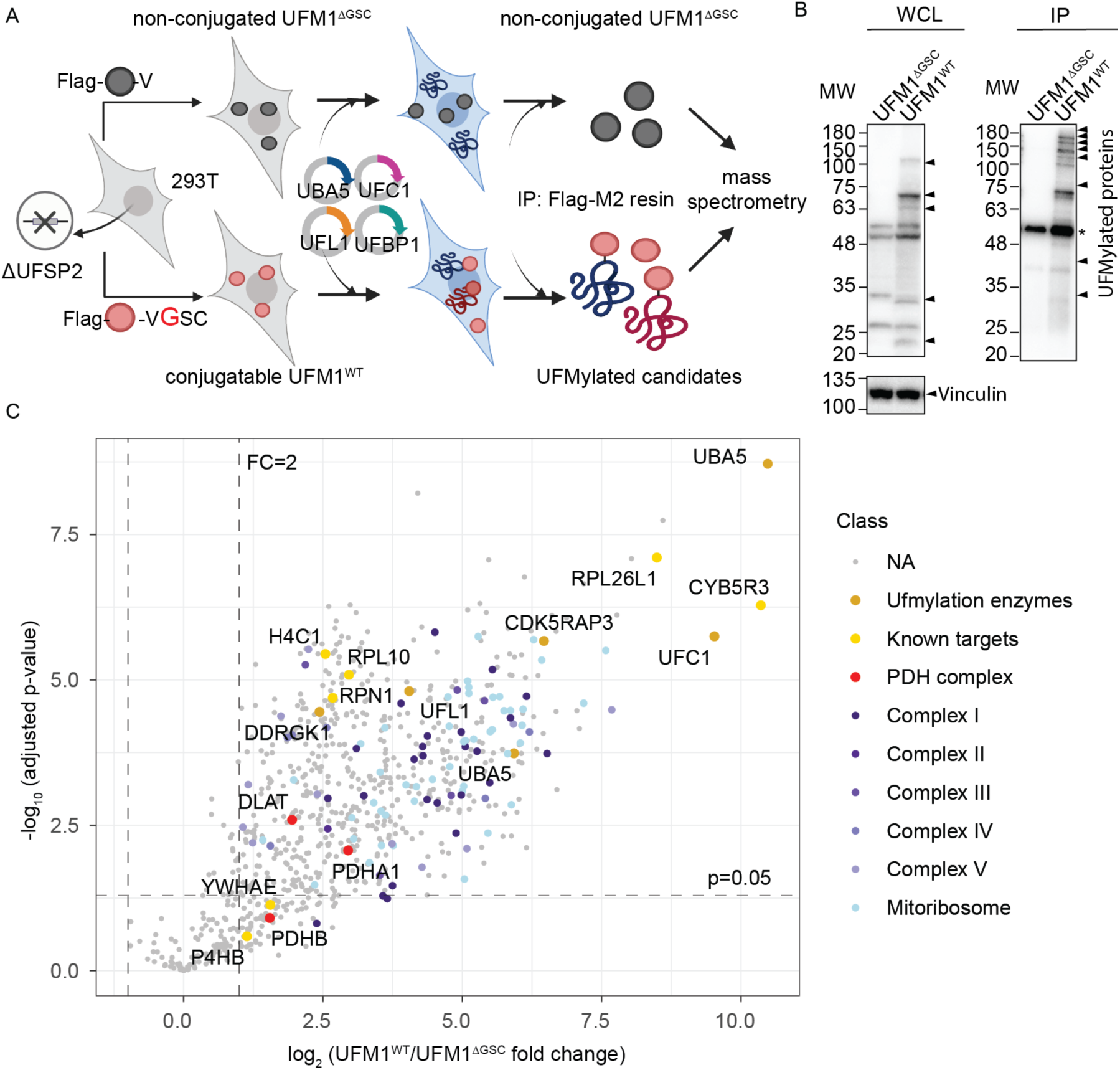
UFM1 immunoprecipitation yields many mitochondrial proteins in UFSP2-deficient cells. **A**: Schematic illustrating an immunoprecipitation-coupled mass spectrometry (IP-MS)-based approach to identify potential UFMylated proteins in cells lacking UFSP2 (created with Biorender.com). **B**: Immunoblotting validating enriched UFMylated candidates in UFM1^WT^ expressing cells compared to UFM1^ΔGSC^-expressing cells. UFMylated proteins were detected using anti-UFM1 antibody. *: non-specific band. **C:** Volcano plot showing enriched UFMylated protein candidates in UFM1^WT^ cells (n=4) compared to UFM1^ΔGSC^ cells (n=4). **Data representation**: Data are presented as the means of n=4 independent biological experiments. **Statistical analysis**: An unpaired, two-tailed Student’s t-test was used for panel (**C**). The statistical cutoff for significance (p<0.05) and the fold change (FC) boundary are indicated.

### UFSP2 ablation increases mitochondrial respiration

To examine the impact of UFMylation on mitochondrial function, we generated isogenic *UFM1*-knockout (ΔUFM1) and *UFSP2*-knockout (ΔUFSP2) HeLa (Fig. 2A-D), HCT116 (Extended Fig. 2A-D) and PANC-1 (Extended Fig. 2E-H) cell lines and measured their oxygen consumption rates (OCR). As expected, *Δ*UFM1 cells lacked detectable UFMylation, whereas *Δ*UFSP2 cells exhibited excessive UFMylation relative to non-targeting control (Control) cells (Fig. 2A, Extended Fig. 2A,E).

**Fig. 2:**
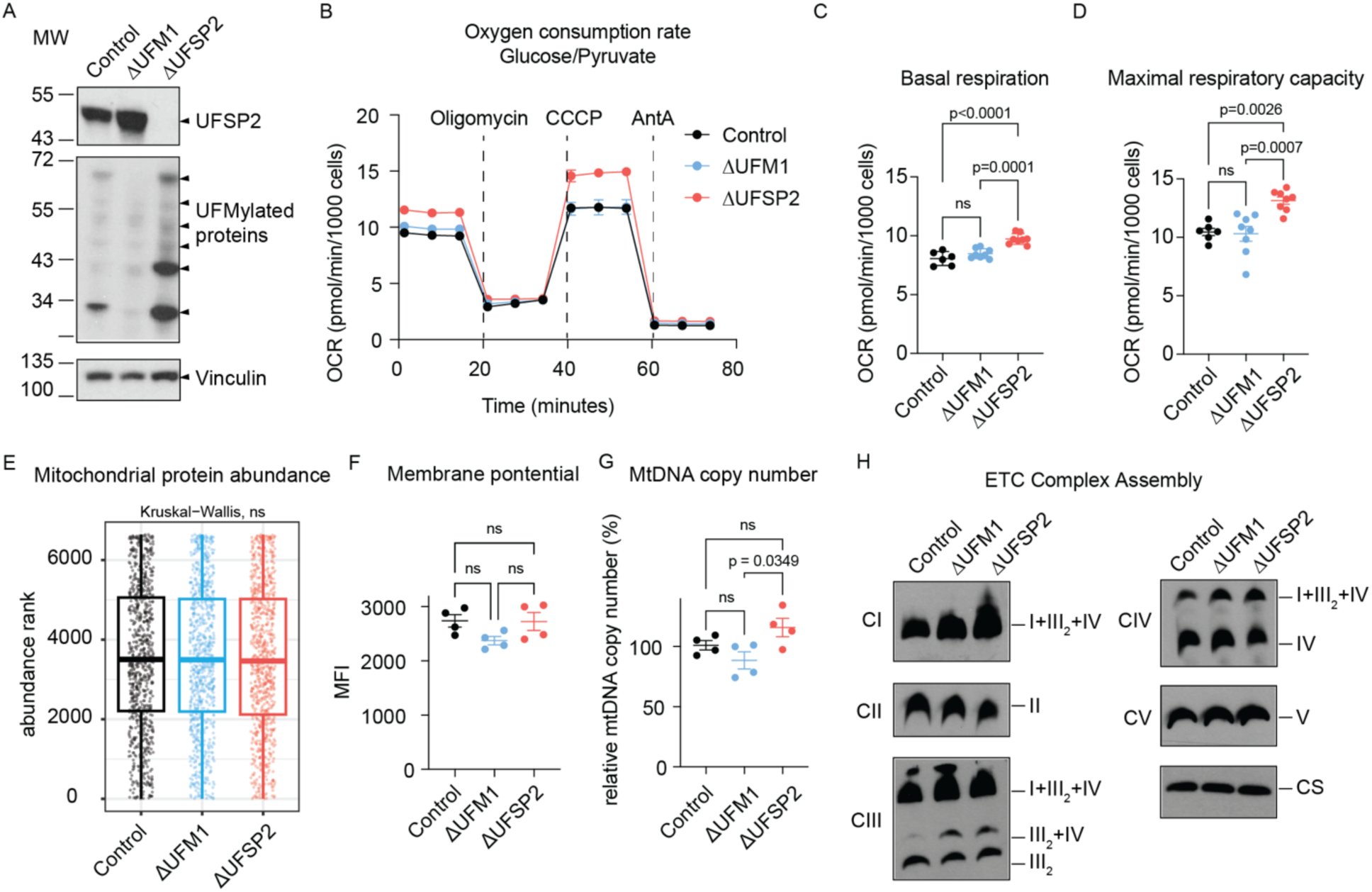
UFSP2 ablation increases mitochondrial respiration. **A**: Western blot validating the effects of UFM1 and UFSP2 knockout on global protein UFMylation. Vinculin serves as a loading control. **B-D**: Oxygen consumption rates (OCR) (**B**), basal respiration (**C**) and maximal respiratory capacity (**D**) of Control (n=6), ΔUFM1 (n=8) and ΔUFSP2 (n=8) HeLa cells. CCCP: Carbonyl cyanide m-chlorophenyl hydrazone, AntA: Antimycin A. **E**: mitochondrial proteins (dots) within each genotype ranked by their median abundance (refer to Methods for details). **F-H**: mitochondrial membrane potential (**F**), mtDNA copy number (**G**), and ETC supercomplexes levels (**H**) in Control (n=4), ΔUFM1 (n=4) and ΔUFSP2 (n=4) HeLa cells. Citrate synthetase (CS) serves as the loading control for supercomplexes. CI – CV: ETC complexes I - V. I+III_2_+IV: supercomplex at 1:2:1 ratio. III_2_+IV: supercomplex at 2:1 ratio. III_2_: complex III dimer. **Data representation**: Mean values are shown in panels **B, E**. In panels **C, D, F, G**, individual data points from independent biological experiments (n) are shown. Horizontal lines and error bars represent the mean ± SEM. **Statistical analysis**: Two-way ANOVA was used for panels **C, D, F, G**. Kruskal-Wallis nonparametric tests were used for panel **E**.

Functionally, while *Δ*UFM1 cells showed no changes in OCR, *Δ*UFSP2 cells exhibited substantially higher basal respiration and maximal respiratory capacity across all three cell lines: HeLa (Fig. 2B-D), HCT116 (Extended Fig. 2B-D), and PANC-1 (Extended Fig. 2F-H) cells. This metabolic shift occurred independently of changes in overall abundance of mitochondrial proteins (Fig. 2E), mitochondrial membrane potential (Fig. 2F), or mitochondrial DNA copy number (Fig. 2G) between control and knockout cells. Furthermore, blue native PAGE analysis revealed no major changes in the assembly or abundance of ETC supercomplexes (Fig. 2H), suggesting that the elevated respiration in ΔUFSP2 cells is driven by enhanced substrate flux rather than structural changes to the respiratory machinery.

### UFSP2 ablation enhances TCA cycle activity via glucose oxidation

To assess changes in TCA cycle activity, we performed time-course stable isotope tracing using uniformly ^13^C-labeled glucose ([U-^13^C]glucose) (Fig. 3A) in Control, *Δ*UFM1 and *Δ*UFSP2 HeLa cells. The fractional enrichments of intracellular glucose m+6 (Extended Fig. 3A), lactate m+3 (Extended Fig. 3B), and pyruvate m+3 (Fig. 3B) were similar in all three cell lines, suggesting that glycolysis is unaffected by the UFMylation status. However, within the first 30 minutes of tracing, *Δ*UFSP2 cells exhibited higher fractional enrichment of citrate m+2 compared to both control and *Δ*UFM1 cells (Fig. 3C). During isotope tracing with [U-^13^C]glucose, citrate m+2 initially comes from the conversion of [U-^13^C]glucose to [U-^13^C]pyruvate via glycolysis, followed by conversion of [U-^13^C]pyruvate to [1,2-^13^C]acetyl-CoA by PDH, and entry of this labeled acetyl-CoA into the TCA cycle. M+2 isotopologues of TCA cycle intermediates and related metabolites downstream of citrate m+2, including alpha-ketoglutarate (*α*-KG), glutamate, succinate, fumarate, malate and aspartate were also elevated in *Δ*UFSP2 cells compared to both Control and *Δ*UFM1 cells (Extended Fig. 3C-H).

**Figure 3:**
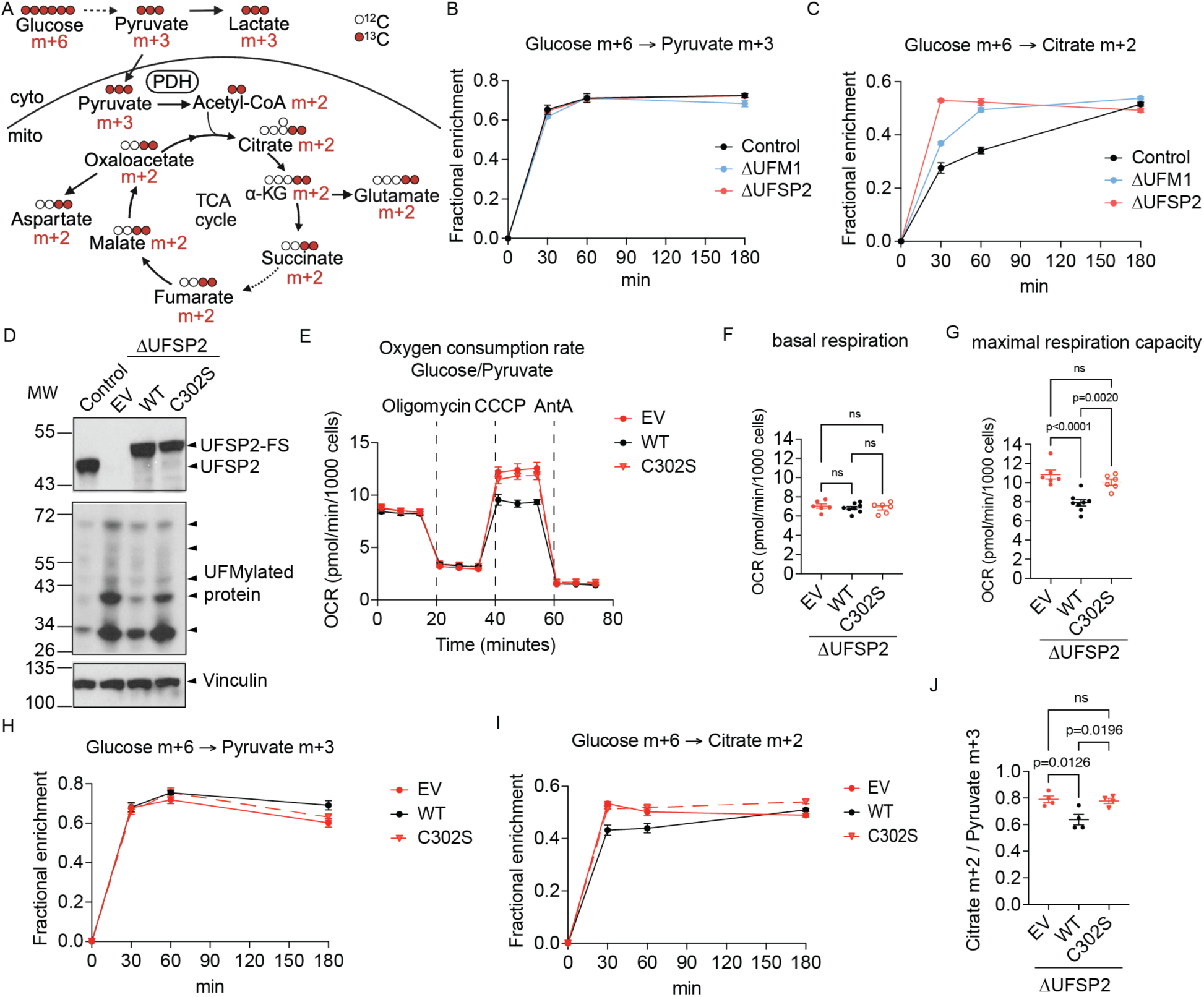
UFSP2 ablation increases labeling of TCA cycle intermediates from [U-^13^C]glucose. **A**: Schematic illustrating stable isotope tracing uniformly ^13^C-labeled glucose ([U-^13^C]glucose) (created with Biorender.com). Cyto: cytoplasm; mito: mitochondrial matrix. **B-C**: Time-dependent fractional enrichment of pyruvate m+3 (B) and citrate m+2 (C) from [U-^13^C]glucose in Control (n=4), ΔUFM1 (n=4), and ΔUFSP2 (n=4) HeLa cells. **D**: Western blot confirming UFSP2 expression and global UFMylation levels in ΔUFSP2 cells ectopically expressing empty vector (EV), Flag-Strep (FS)-tagged wildtype (WT) UFSP2, or the catalytically inactive C302S UFSP2 variant. **E-G**: oxygen consumption rates (OCR) (**E**), basal respiration (**F**) and maximal respiratory capacity (**G**) of the isogenic lines described in (**D**) (EV (n=6), WT (n=8), and C302S (n=6) UFSP2). CCCP: Carbonyl cyanide m-chlorophenyl hydrazone, AntA: Antimycin A. **H-J**: Time-dependent fractional enrichment of pyruvate m+3 (**H**) and citrate m+2 (**I**) from [U-^13^C]glucose and citrate m+2 to pyruvate m+3 ratio (**J**) in ΔUFSP2 cells ectopically expressing EV (n=4), WT (n=4), and C302S (n=4) UFSP2. **Data representation**: Mean values are shown in panels **B, C, E, H, I**. Individual data points from the indicated number of independent biological experiments (n) are shown in panels **F, G, J**. Horizontal lines and error bars represent the mean ± SEM. **Statistical analysis**: Two-Way ANOVA was used for statistical analyses in panels **F, G, J**.

To confirm that this phenotype specifically results from the loss of UFSP2 enzymatic activity, we reconstituted *Δ*UFSP2 HeLa cells with an empty vector (EV), wildtype UFSP2 (WT), or a catalytically dead UFSP2 mutant (C302S). As expected, the EV and the C302S variant failed to clear accumulated UFMylated substrates compared to re-expressing WT UFSP2 in *Δ*UFSP2 HeLa cells (Fig. 3D). Oxygen consumption analyses (Fig. 3E) showed comparable basal respiratory rates (Fig. 3F), but higher maximal respiratory capacity (Fig. 3G) in *Δ*UFSP2 cells expressing either EV or C302S variant compared to WT UFSP2. We then repeated the [U-^13^C]glucose tracing experiment in these rescued lines. While labeling of glucose (Extended Fig. 4A), lactate (Extended Fig. 4B) and pyruvate (Fig. 3H) were similar in all three cell lines, reconstitution with WT UFSP2 reduced citrate m+2 labeling at early time points, but reconstitution with C302S UFSP2 did not have this effect (Fig. 3I). Labeling in other TCA cycle intermediates followed a similar pattern (Extended Fig. 4C-H).

Notably, glutamine-mediated anaplerosis was unaffected by the lack of UFSP2 as evidenced by the comparable fractional enrichment of labeled TCA cycle intermediates from uniformly ^13^C-labeled glutamine ([U-^13^C]glutamine) tracing (Extended Fig. 5). Collectively, these data suggest that UFSP2 controls glucose metabolism upstream of the TCA cycle. In line with this, compared to cells expressing WT UFSP2, the higher citrate m+2/pyruvate m+3 ratios in cells lacking enzymatically functional UFSP2 suggests that UFSP2 regulates glucose oxidation at the level of PDH (Fig. 3J).

### UFSP2 ablation enhances PDH enzyme activity

To assess PDH function, we next examined the fractional enrichment of acetyl-CoA m+2 derived from [U-^13^C]glucose (schematic illustration in Fig. 3A). During culture with [U-^13^C]glucose, HeLa cells reconstituted with either the EV or the C302S variant exhibited a marked, time-dependent increase in acetyl-CoA m+2 fractional enrichment compared to cells expressing WT UFSP2 (Fig. 4A-B). Notably, the total cellular acetyl-CoA pools remained comparable across all three cell lines (Fig. 4C), suggesting that the difference in labeling reflects an increase in the rate of *de novo* synthesis via the PDH complex rather than changes in the steady-state pool size.

**Fig. 4:**
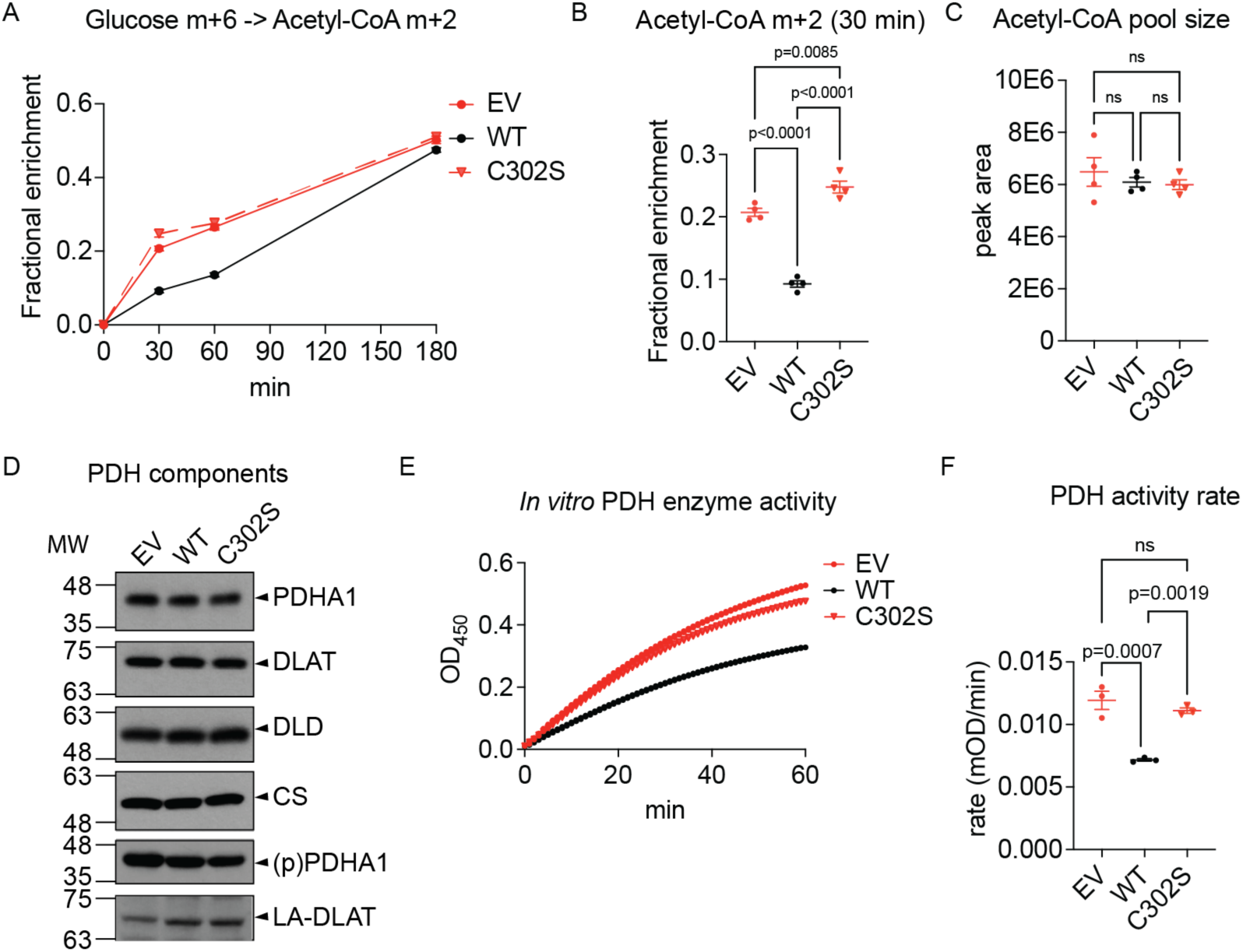
UFSP2 enzyme activity regulates the PDH complex. **A-B**: Fractional enrichment of acetyl-CoA m+2 over time (**A**) and after 30 minutes of culture (**B**) supplemented with [U-^13^C]glucose in ΔUFSP2 cells ectopically expressing EV (n=4), WT (n=4) or C302S (n=4) UFSP2. **C**: Relative steady-state acetyl-CoA abundance in the indicated cell lines (n=4). **D-F**: Protein abundance, phosphorylation and lipoylation status of PDH components. Citrate synthase (CS) serves as a loading control (**D**), *in vitro* PDH enzyme activity (**E**) and rate (**F**) in ΔUFSP2 cells ectopically expressing EV (n=3), WT (n=3) or C302S (n=3) UFSP2. **Data representation**: Mean values are shown in panels **A, E**. Individual data points from the indicated number of independent biological experiments (n) are shown in panels **B, C, F**. Horizontal lines and error bars represent the mean ± SEM. **Statistical analysis**: Two-Way ANOVA was used for statistical analyses in panels **B, C, F**.

In principle, increased acetyl-CoA synthesis via the PDH complex could result from either enhanced PDH protein abundance or enhanced enzymatic activity. However, we observed no changes in the protein abundance of the PDH subunits, PDHA1, DLAT, and DLD, among isogenic ΔUFSP2 cells expressing EV, WT, and C302S UFSP2 (Fig. 4D). Therefore, we measured PDH enzymatic activity directly using an immunoprecipitation-based activity assay. Consistent with our isotope tracing results, the expression of EV or C302S variant resulted in increased PDH activity (Fig. 4E-F) compared to cells expressing WT UFSP2. These data indicate that USFP2 regulates the entry of glucose-derived carbon into the TCA cycle by modulation of PDH catalytic activity.

If the enhanced oxidative phosphorylation observed in *Δ*UFSP2 cells is mediated by increased PDH activity, we reasoned that providing an alternative, PDH-independent source of acetyl-CoA would normalize respiration. To test this, we supplemented cells with acetate (Extended Fig. 6A), which can be converted to acetyl-CoA by acetyl-CoA synthetase (ACSS1)^51^. Indeed, UFSP2 ablation no longer boosted respiration in cells fed with acetate instead of glucose and pyruvate (Extended Fig. 6B-D). This suggests that UFMylation regulates mitochondrial respiration specifically at the PDH-mediated pyruvate oxidation step.

Interestingly, we did not detect differences in the inhibitory phosphorylation of PDHA1 or the activating lipoylation of DLAT (Fig. 4D), both of which are canonical regulators of PDH activity^52–55^. This absence of known regulatory changes raises the possibility that UFMylation itself regulates PDH function.

### UFMylation of DLAT increases PDH enzyme activity

IP-MS identified several components of the PDH complex, including PDHA1, PDHB and DLAT as potential UFMylation target components (Fig. 1C). However, it has been suggested that overexpression of the E1-E2-E3 enzyme components of the UFMylation pathway can lead to non-physiologically UFMylated artifacts in IP-MS assays^19,56^. To address this, we designed a secondary validation assay for UFMylated candidates in UFSP2-KO HEK293F cells expressing either UFM1^WT^ or UFM1^ΔGSC^ (Fig. 5A). We transiently expressed Flag-Strep-tagged PDHA1 (Extended Fig. 7A), PDHB (Extended Fig. 7B), DLD (Extended Fig. 7C) and DLAT (Fig. 5A) in these isogenic cell lines. Immunoblotting following affinity pulldown confirmed that DLAT, an E2 component of the PDH complex, is UFMylated, detected by a shift in its molecular weight by UFM1 addition and confirmed by immunoblotting using an anti-UFM1 antibody (Fig. 5B). Subsequent LC/MS-MS analysis of the DLAT tryptic digest identified three potential lysine residues as UFMylation sites: K118 (Extended Fig. 8A), K363 (Extended Fig. 8B), and K547 (Extended Fig. 8C). For the other subunits of the PDH complex, the anti-UFM1 antibody detected a faint signal for PDHB, and no signal for the others (Extended Fig. 7A-C).

**Fig. 5:**
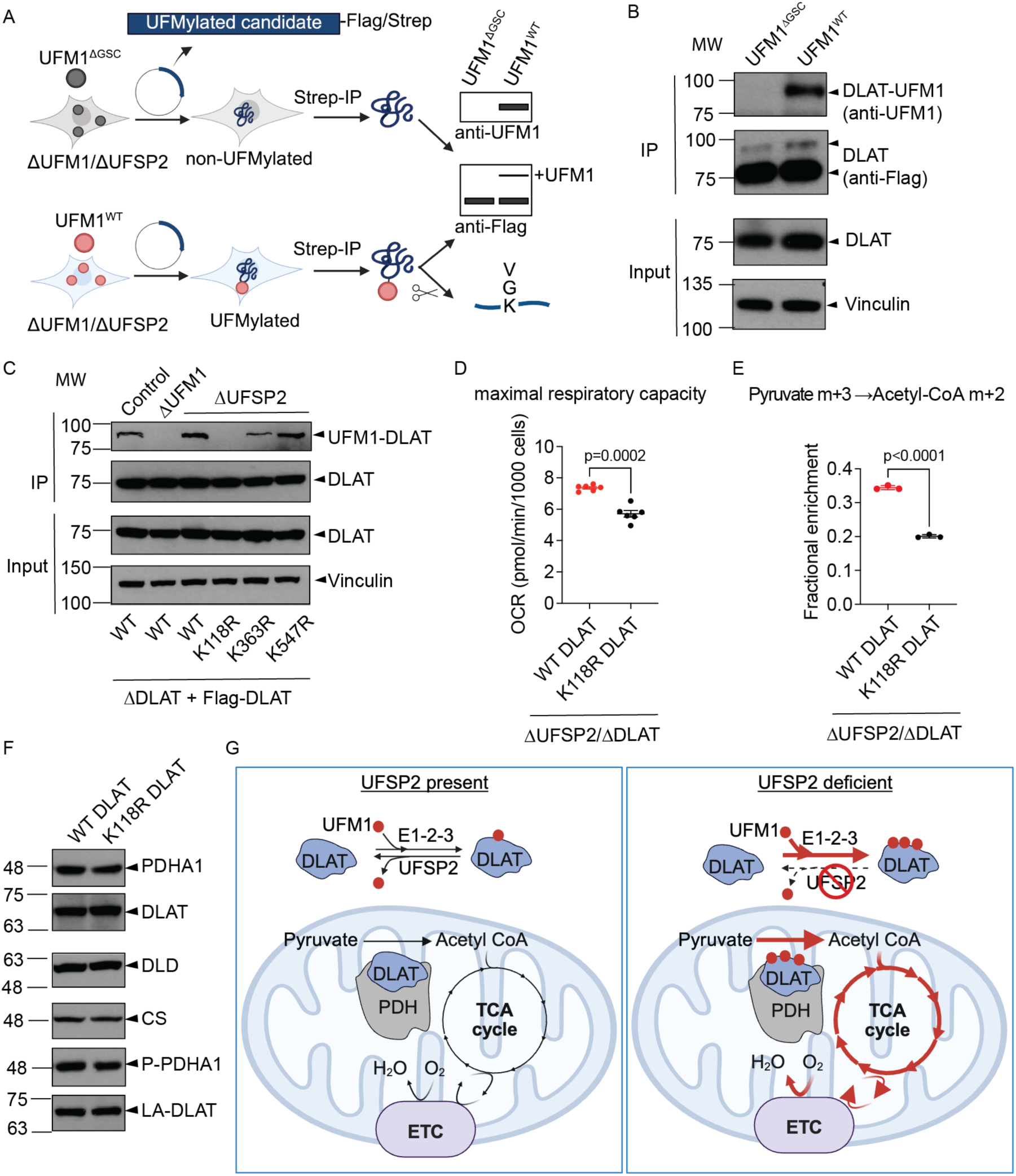
DLAT UFMylation regulates PDH enzyme activity. **A**: Schematic of in-cell UFMylation assays used to validate UFMylation targets (created with Biorender.com). **B**: Validation of DLAT as a bona fide UFMylated target (**B**). **C**: K118R mutation abolishes DLAT UFMylation. **D-F**: Maximal respiratory capacity (**D**), fractional enrichment of acetyl-CoA m+2 from [U-^13^C]pyruvate m+3 (**E**), protein abundance, phosphorylation and lipoylation of PDH components in ΔUFSP2/ΔDLAT cells reconstituted with either WT or K118R DLAT (**F**). Citrate synthase (CS) serves as a loading control. P-PDHA1: phosphorylated PDHA1. LA-DLAT: lipoylated DLAT. **G**: Model illustrating how UFMylation of the PDH complex regulates pyruvate oxidation in mitochondria. **Data representation**: Individual data points from the indicated number of independent biological experiments (n) are shown in panels **D, E**. Horizontal lines and error bars represent the mean ± SEM. **Statistical analysis**: An unpaired, two-sided Student’s t-test was used for statistical analyses in panels **D, E**.

To validate DLAT’s UFMylated sites, we generated *DLAT* knockout (ΔDLAT) HeLa cells (Extended Fig. 9) and reconstituted them with either Flag-Strep-tagged WT DLAT or DLAT alleles with mutations at the relevant lysine residues (K118R, K363R, or K547R). Following affinity purification of DLAT, we assessed its UFMylation status. Consistent with our previous findings, WT DLAT had higher UFMylation levels in *Δ*UFSP2 cells compared to control cells, whereas no UFMylation was detected in *Δ*UFM1 cells (Fig. 5C). Among the three lysine-to-arginine mutations, the K118R mutation had the greatest effect on DLAT UFMylation. In contrast, the K363R mutation had only a modest effect, and K547R had virtually no impact on UFMylation levels (Fig. 5C).

Functionally, the expression of K118R DLAT in *Δ*UFSP2/ΔDLAT cells reduced both basal respiration and maximal respiration compared to the re-expression of WT DLAT (Fig. 5D, Extended Fig. 10). To further confirm the metabolic impact, we performed stable isotope tracing using uniformly ^13^C-labeled pyruvate ([U-^13^C] pyruvate m+3). The fractional enrichment of acetyl-CoA m+2 from [U-^13^C] Pyruvate m+3 was lower in *Δ*UFSP2 cells expressing K118R DLAT (Fig. 5E), despite no changes in total PDH protein abundance in WT and K118R DLAT cells (Fig. 5F). Collectively, these results demonstrate that UFMylation of DLAT specifically at K118 increases enzymatic activity of the PDH complex (Fig. 5G).

## DISCUSSION

The discovery of the UFMylation pathway over two decades ago redefined our understanding of ubiquitin-like (UBL) modifications^1^, yet its functional landscape has remained largely confined to the endoplasmic reticulum^20–24^ and nucleus^26–28^. Our study expands this paradigm by demonstrating that loss of UFSP2/deUFMylation activity, which is observed in cells derived from patients with pathogenic UFSP2 mutations^41–43^, results in the aberrant UFMylation of mitochondrial proteins. By identifying the dihydrolipoamide S-acetyltransferase (DLAT) subunit of the pyruvate dehydrogenase (PDH) complex as a substrate that hyper-accumulates UFM1 tags in the absence of UFSP2, we provide a potential biochemical dimension of UFSP2-related disorders.

The PDH complex serves as the critical metabolic bridge between glycolysis and the TCA cycle, requiring the precise assembly of E1 (PDH), E2 (DLAT) and E3 (DLD) subunits. While inhibitory phosphorylation of the E1 subunit by PDH kinases is a well-characterized “brake” on PDH activity^52^, our finding that UFMylation at K118 on the E2 (DLAT) subunit activates PDH activity introduces a previously unrecognized layer of positive regulation. We suggest that in patients harboring UFSP2 variants, such as V115E (linked to neurodevelopmental disorders) or Y280H, D426A, and H428R (linked to skeletal dysplasias), the inability to remove UFM1 from DLAT K118 creates a state of unregulated mitochondrial hyperactivity.

Our observation that UFSP2 deletion enhances maximal respiration while respiration in UFM1-deleted or control cells is essentially unchanged suggests that mitochondria maintain a “UFMylation threshold”. Under homeostatic conditions, UFSP2-mediated de-UFMylation keeps PDH activity in check; however, the loss of UFSP2 tips this balance, leading to an activation of PDH activity. Mechanistically, we propose that UFM1 conjugation at K118 acts as a gain-of-function modification. K118 is located immediately adjacent to the K132 lipoylation site within the lipoyl domain, which acts as a “swinging arm” to transfer the acetyl group from E1 to Coenzyme A (CoA) to form acetyl-CoA^54^. We speculate that UFM1 conjugation induces steric or electrostatic shifts that enhance the conformational flexibility of the lipoyl arm. This increased mobility may accelerate the transfer of acetyl groups, lowering the energetic barrier for substrate channeling and increasing acetyl-CoA availability to fuel the TCA cycle (Fig. 5G).

The link between UFSP2 deficiency and mitochondrial hyperactivity provides a new perspective on UFSP2-related human disease. UFSP2 deficiency has been observed in multiple cancer types^57^ and in the brains of patients with Alzheimer’s dementia^58^, underscoring a potentially broad role in human pathology. In long-lived post-mitotic cells like neurons, loss of the UFSP2-mediated metabolic brake could trigger chronic oxidative stress via the overproduction of reactive oxygen species (ROS), resulting in mitochondrial damage, including mitochondrial DNA mutations, mitochondrial membrane permeability, and perturbed Ca^2+^ homeostasis^59,60^. This could contribute to the synaptic loss and neurodegeneration characteristic of AD^59,60^.

Our discovery of the UFSP2-PDH axis reveals a previously underappreciated function of the UFMylation pathway in regulating mitochondrial metabolism. While this study focuses specifically on the PDH complex, our proteomic data uncovered a potential network of UFMylation targets within the core mitochondrial machinery, including mitochondrial ribosomal proteins (MRPs) and subunits of the electron transport chain (ETC) Complexes I - V (Fig. 1C). These findings suggest that UFMylation may exert multi-layered control over mitochondrial bioenergetics extending beyond pyruvate oxidation. Future exploration of these additional mitochondrial substrates will further define how the UFMylation/deUFMylation balance serves as a switch for mitochondrial metabolism in both health and disease.

---

**Extended Fig. 1:**
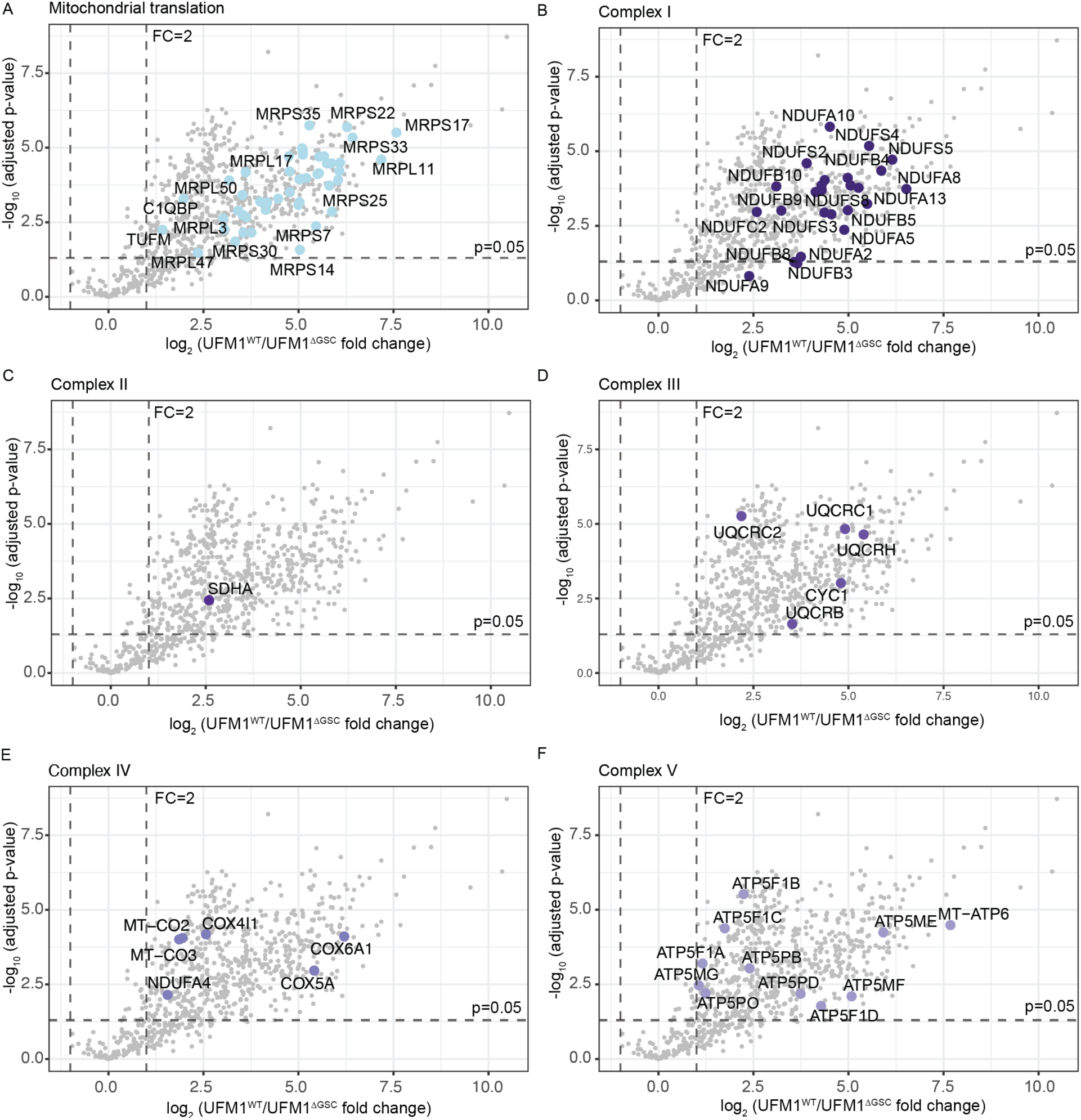
Components of the mitochondrial translation machinery and the ETC complexes are enriched among UFMylated targets. A-F: Volcano plots highlighting potentially UFMylated proteins involved in mitochondrial translation (A) and ETC complexes I-V (B-F). Data representation: Data are presented as the means of n=4 independent biological experiments. Statistical analysis: An unpaired, two-tailed Student’s t-test was used for panels A-F. The statistical cutoff for significance (p<0.05) and the fold change (FC) boundary are indicated.

**Extended Fig. 2:**
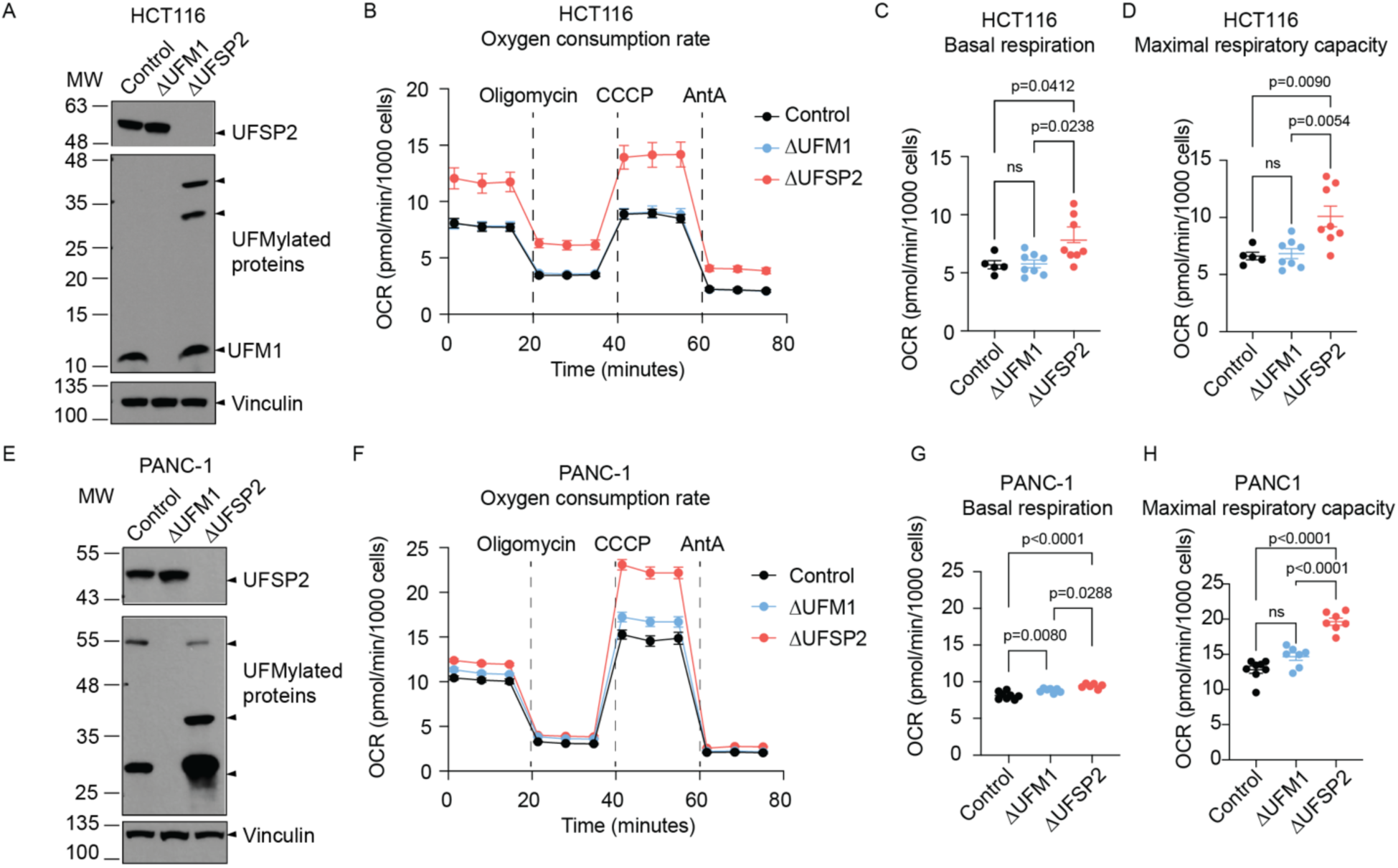
UFSP2 ablation increases mitochondrial respiration across multiple cancer cell lines. **A-D**: Analysis of HCT116 cells. **A**: Western blot validating the effects of UFM1 and UFSP2 knockout on global protein UFMylation. Vinculin serves as the loading control. **B-D**: Oxygen consumption rates (OCR) (**B**), basal respiration (**C**) and maximal respiratory capacity (**D**) in control (n=5), ΔUFM1 (n=8) and ΔUFSP2 (n=8) cells. **E-H**: Analysis of PANC-1 cells. **E**: Western blot validating the effects of UFM1 and UFSP2 knockout on global protein UFMylation. **F-H**: Oxygen consumption rates (OCR) (**F**), basal respiration (**G**) and maximal respiratory capacity (**H**) in Control (n=8), ΔUFM1 (n=7) and ΔUFSP2 (n=7) PANC-1 cells. CCCP: Carbonyl cyanide m-chlorophenyl hydrazone, AntA: Antimycin A. **Data representation**: Mean OCR values are shown in panels **B, F**. In panels **C, D, G, H**, individual data points from the indicated number of independent biological experiments (n) are shown. Horizontal lines and error bars represent the mean ± SEM. **Statistical analysis**: Two-way ANOVA was used for panels **C, D, G, H**.

**Extended Fig. 3:**
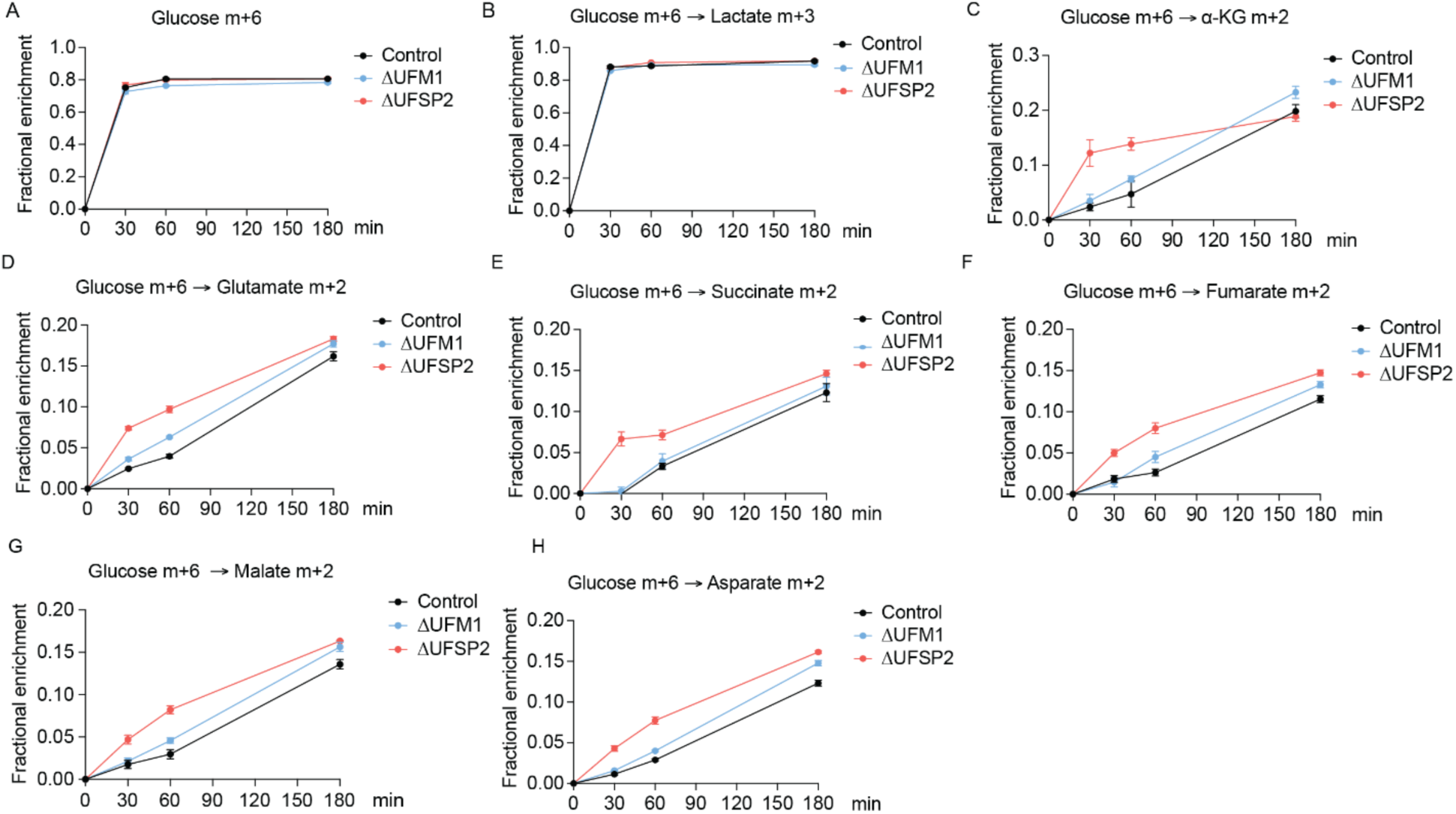
UFSP2 ablation increases labeling of TCA cycle intermediates from [U-^13^C]glucose. Time-dependent fractional enrichment of glucose m+6 (**A**), lactate m+3 (**B**), alpha-ketoglutarate (*α*-KG) m+2 (**C**), glutamate m+2 (**D**), succinate m+2 (**E**), fumarate m+2 (**F**), malate m+2 (**G**), and aspartate m+2 (**H**) from [U-^13^C]glucose in Control (n=4), ΔUFM1 (n=4), and ΔUFSP2 (n=4) cells. **Data representation**: all data are presented as the mean ± SEM of n=4 independent biological experiments.

**Extended Fig. 4:**
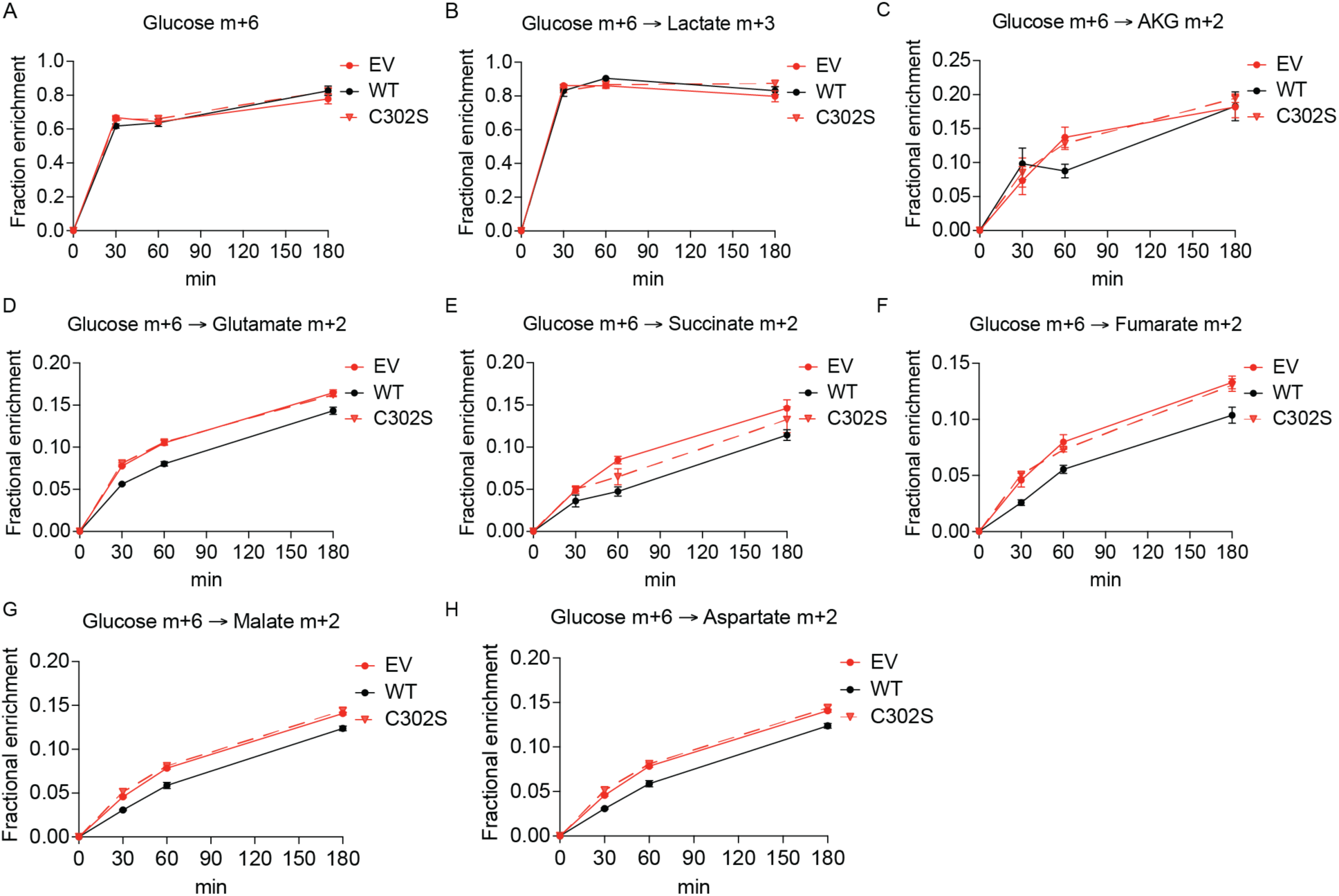
UFSP2 enzyme activity regulates labeling of TCA cycle intermediate. Time-dependent fractional enrichment of glucose m+6 (**A**), lactate m+3 (**B**), alpha-ketoglutarate (*α*-KG) m+2 (**C**), glutamate m+2 (**D**), succinate m+2 (**E**), fumarate m+2 (**F**), malate m+2 (**G**), and aspartate m+2 (**H**) from [U-^13^C]glucose in ΔUFSP2 cells ectopically expressing EV (n=4), WT (n=4) or C302S (n=4) UFSP2. **Data representation**: all data are presented as the mean ± SEM of n=4 independent biological experiments.

**Extended Fig. 5:**
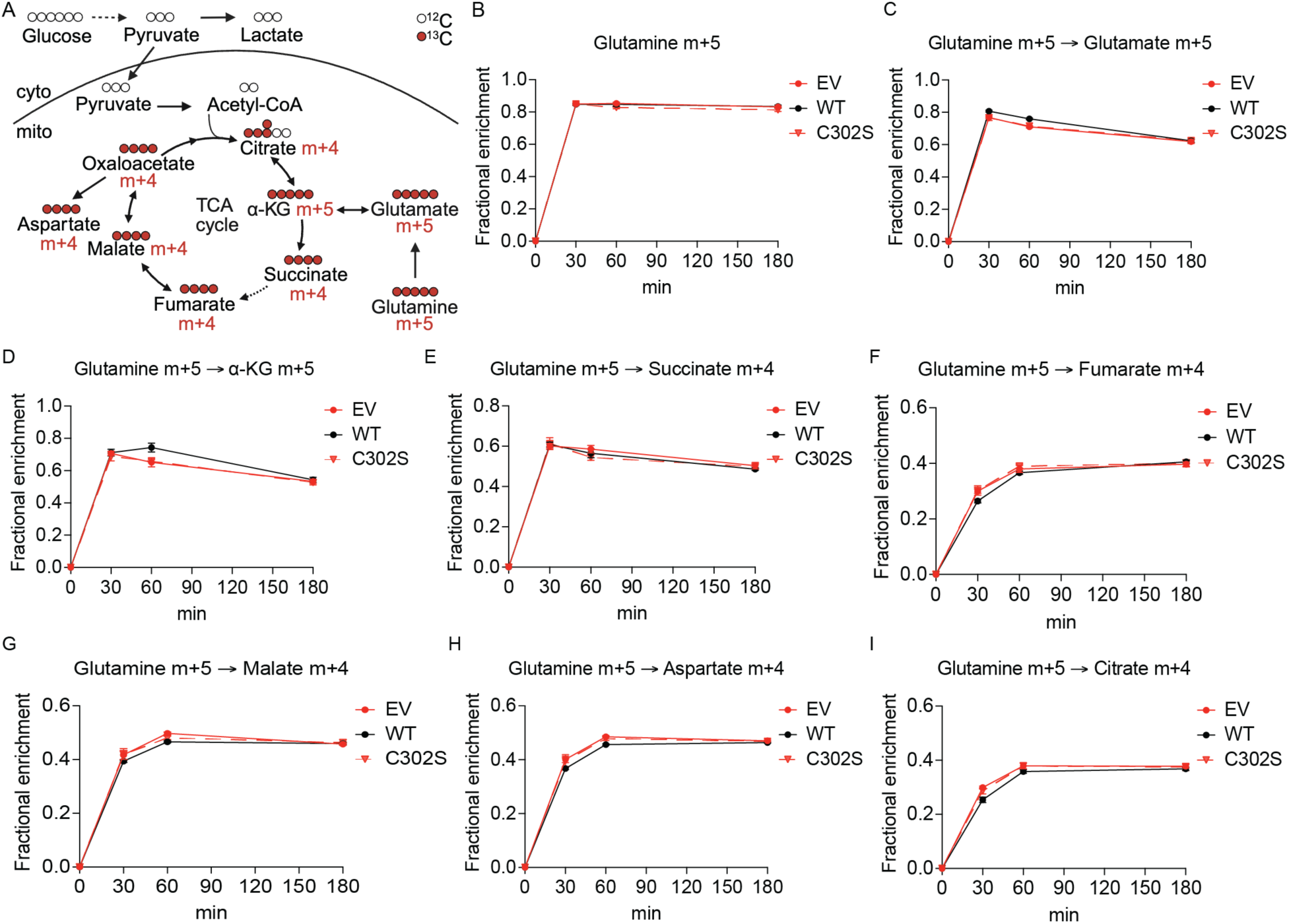
UFSP2 ablation does not change labeling of TCA cycle intermediates from [U-^13^C]glutamine. A: Schematic for uniformly ^13^C-labeled glutamine ([U-^13^C]glutamine) tracing (created with Biorender.com). Cyto: cytoplasm; mito: mitochondrial matrix. B-I: Time-dependent fractional enrichment of glutamine m+5 (B), glutamate m+5 (C), *α*-KG m+5 (D), succinate m+4 (E), fumarate m+4 (F), malate m+4 (G), aspartate m+4 (H), citrate m+4 (I) from [U-^13^C]glutamine in ΔUFSP2 cells ectopically expressing EV (n=4), WT (n=4) or C302S (n=4) UFSP2. **Data representation**: all data are presented as the mean ± SEM of n=4 independent biological experiments.

**Extended Fig. 6:**
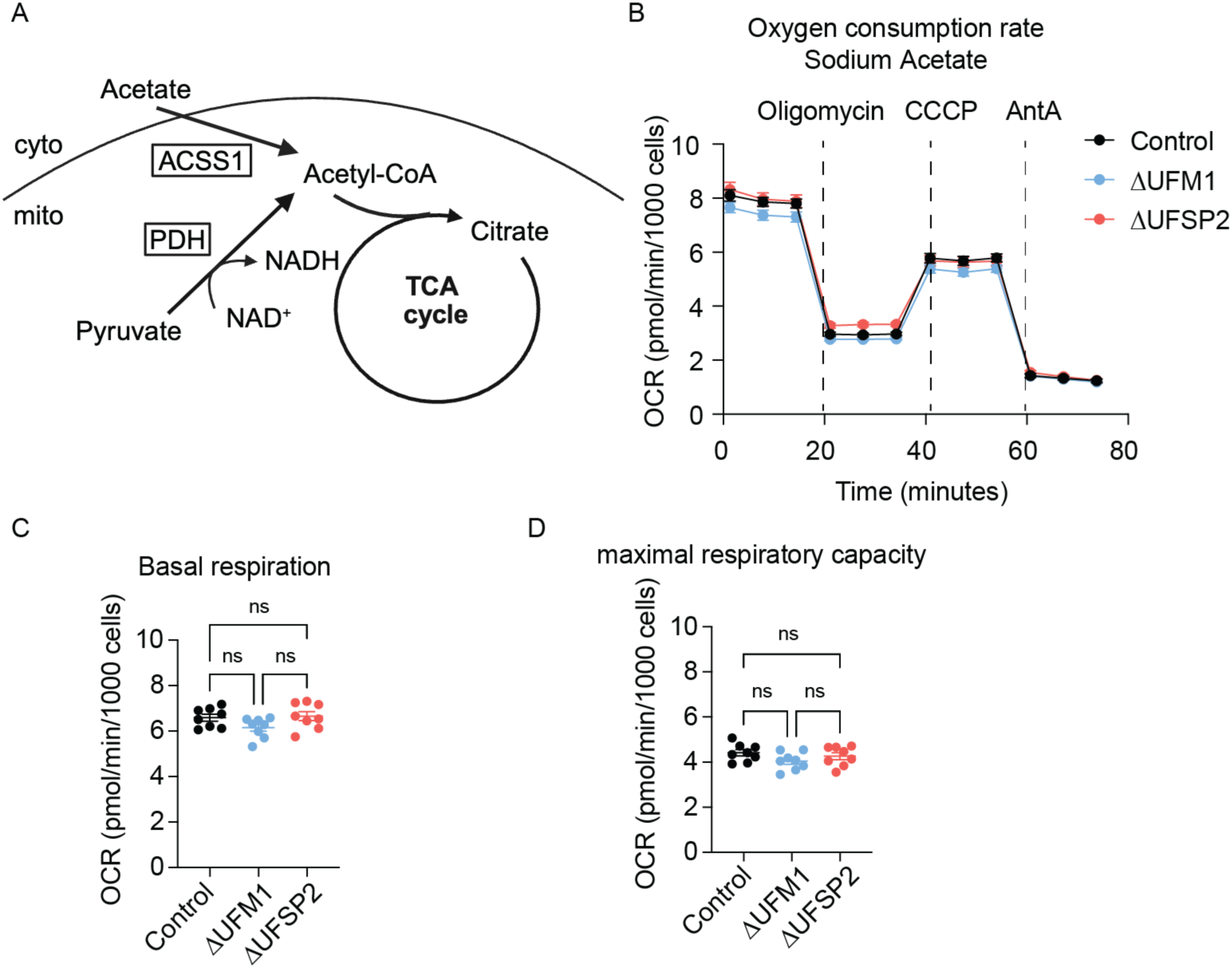
Bypassing PDH-dependent acetyl-CoA production reverses respiratory phenotypes in ΔUFSP2 cells. **A**: schematic illustrating ACSS1-dependent acetyl-CoA production from sodium acetate (created with Biorender.com). Cyto: cytoplasm; mito: mitochondrial matrix. **B-D**: Oxygen consumption rates (OCR) (**B**), basal respiration (**C**) and maximal respiratory capacity (**D**) from Control (n=8), ΔUFM1 (n=8) and ΔUFSP2 (n=8) cells. CCCP: Carbonyl cyanide m-chlorophenyl hydrazone, AntA: Antimycin A. **Data representation**: Mean OCR values are shown in panel **B**. Individual data points from the indicated number of independent biological experiments (n) are shown in panels **C, D**. Horizontal lines and error bars represent the mean ± SEM. **Statistical analysis**: Two-Way ANOVA was used for statistical analyses in panels **C, D**.

**Extended Fig. 7:**
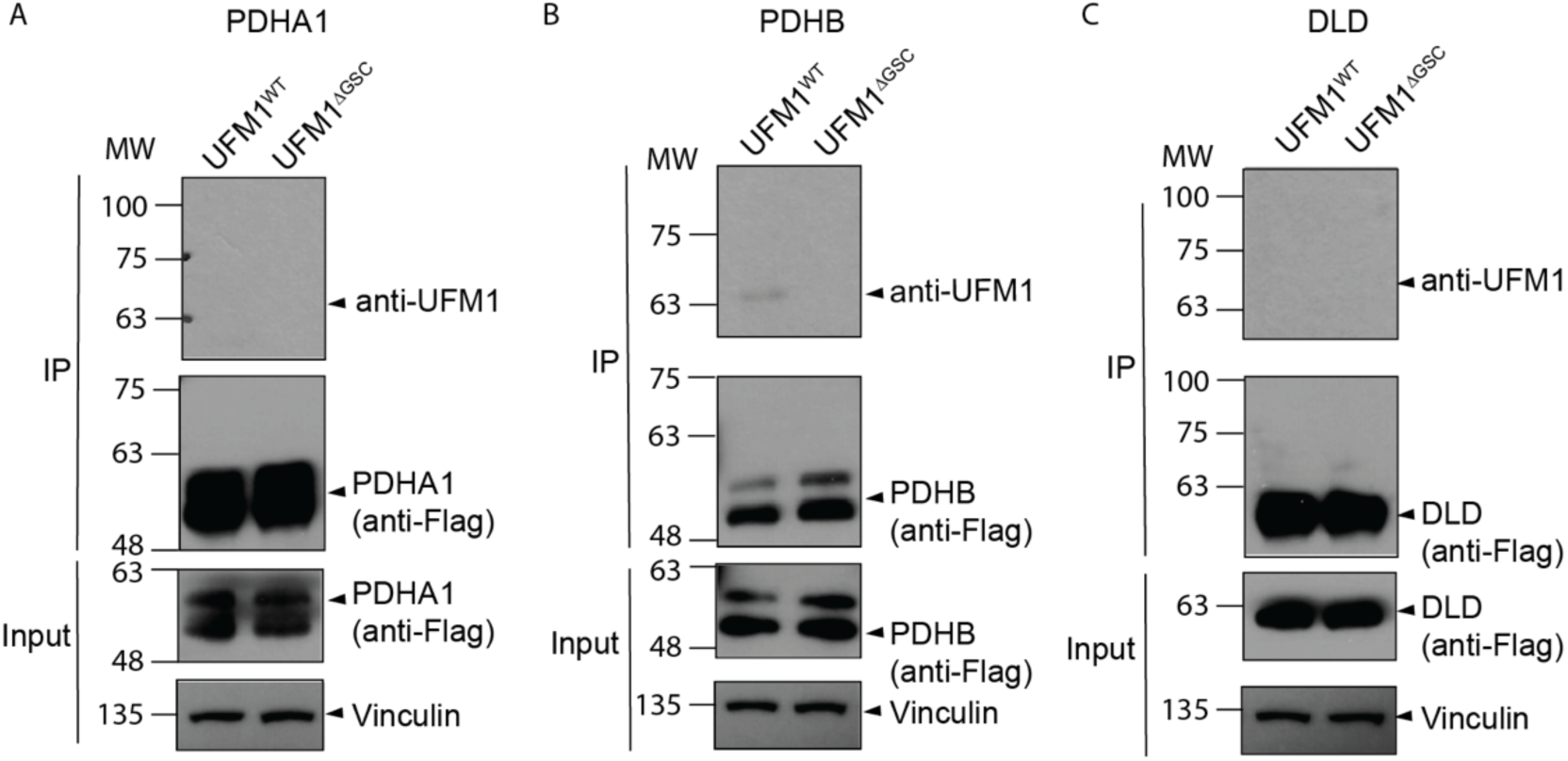
Assessment of components of PDH enzyme as UFMylated targets. validation of PDHA1 (A), PDHB (B) and DLD (C) as a UFMylated target by in-cell UFMylation assays.

**Extended Fig. 8:**
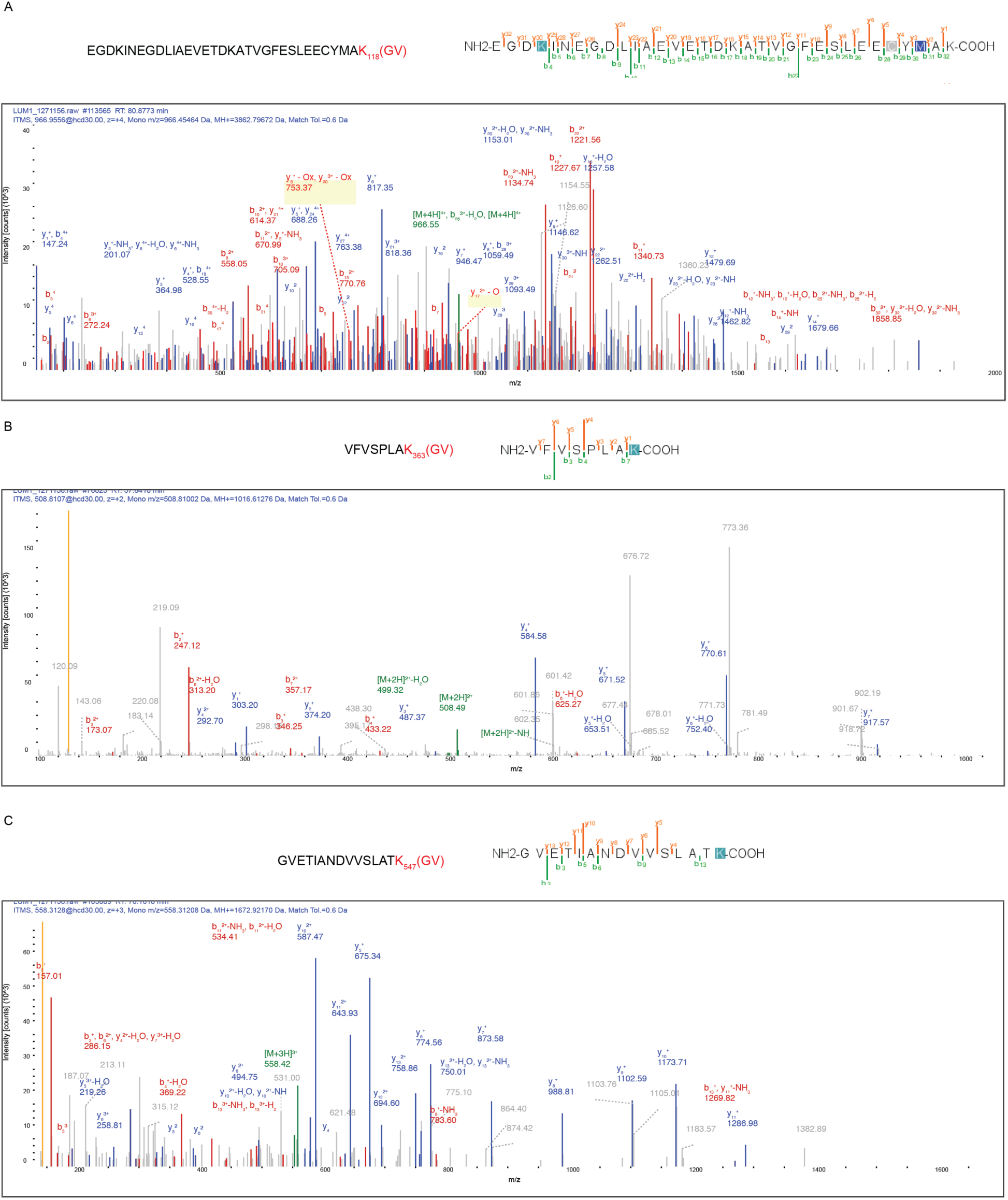
Identification of UFMylated lysine residues on DLAT. LC-MS/MS spectra of the peptides whose lysine residues, K118 (A), K363 (B), and K547 (C) were identified as UFMylation sites.

**Extended Fig. 9:**
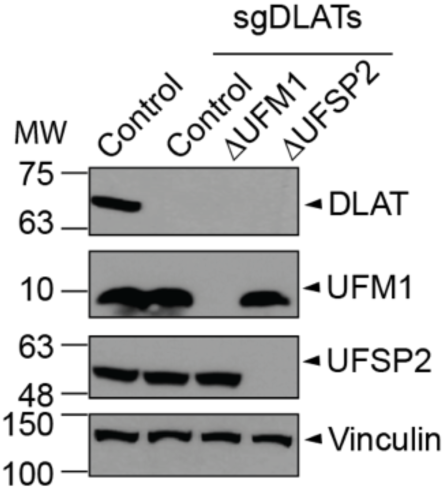
CRISPR-Cas9-mediated knockout of DLAT in HeLa isogenic cell lines. sgDLATs: guide RNAs targeting *DLAT* gene. Vinculin serves as a loading control.

**Extended Fig. 10:**
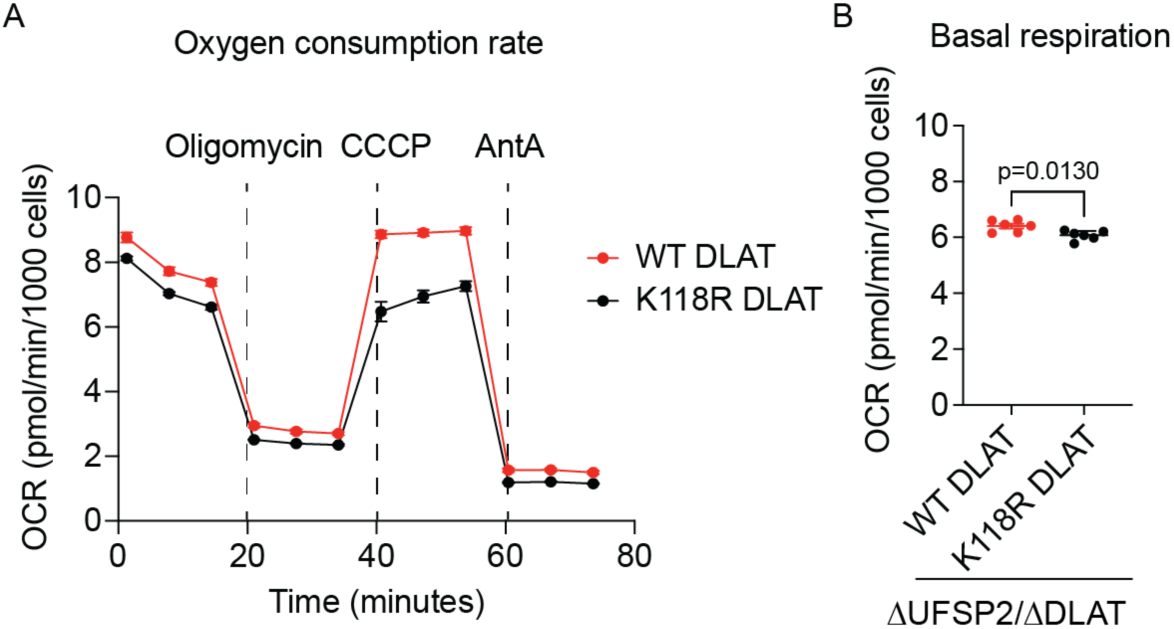
Abolishing UFMylation of DLAT reduces mitochondrial respiration in ΔUFSP2 HeLa cell line. Oxygen consumption rates. (A) and mitochondrial basal respiration (B) of the isogenic ΔUFSP2/ΔDLAT HeLa cells ectopically expressing WT or K118R DLAT. CCCP: Carbonyl cyanide m-chlorophenyl hydrazone, AntA: Antimycin A. **Data representation**: Mean OCR values are shown in panel (**A**). Individual data points from the indicated number of independent biological experiments (n) are shown in panel **B**. Horizontal lines and error bars represent the mean ± SEM. **Statistical analysis**: Two-Way ANOVA was used for statistical analyses in panel **B**.

## Acknowledgments

We thank members of the DeBerardinis laboratory for helpful discussions; the UT Southwestern Proteomics Core and CRI’s Flow Cytometry Core for assistance with proteomics and cell sorting, respectively. This article is subject to HHMI’s Open Access to Publications policy. HHMI lab heads have previously granted a nonexclusive CC BY 4.0 license to the public and a sublicensable license to HHMI in their research articles. Pursuant to those licenses, the author-accepted manuscript of this article can be made freely available under a CC BY4.0 license immediately upon publication.

## Author Contributions

R.J.D. and P.T.N. conceived and designed the study. P.T.N and Z.W. performed most of the experiments and data analyses, assisted by D.K., T.O., S.Y., V.S., T.T., L.C., D.D., M.C., H.L. F.C. F.C. designed, optimized and validated the metabolomics methods. A.J., E.S. and J.J.C. performed mass spectrometry experiments and data analyses. D.K. and G.H. helped with the LC-MS experiments and data analyses. P.M. and M.N. provided intellectual contributions and expert feedback on mitochondrial functional assays and cellular metabolism characterization, respectively. P.T.N. and R.J.D. wrote the paper and all the other authors edited and commented on the paper.

## Methods

### Cell culture

HeLa, HCT-116 and PANC-1 cells were kindly provided by Dr. Gerta Hoxhaj (UT Southwestern). 293T cells were purchased from American Type Culture Collection (ATCC, CRL-3216). These adherent lines were maintained in RPMI-1640 (Thermo Fisher Scientific, CB-40234), supplemented with 10% fetal bovine serum (FBS) and cultured at 37°C in a humidified atmosphere containing 5% CO_2_. HEK 293F suspension cells (Invitrogen, R79007) were grown in FreeStyle^TM^ 293 Expression Medium (Thermo Fisher Scientific, 12-338-026) at 37°C with 8% CO_2_ and constant orbital shaking at 130 rpm. All cell lines were routinely screened for mycoplasma contamination and confirmed negative.

### Gene deletion and overexpression

To generate *UFM1*, *UFSP2, or DLAT* knockouts using the Cas9 D10A nickase system, cells were transfected with two modified pDG461 constructs^61^ with mGreenLantern and mScarlett fluorescent markers. Each vector contains two sgRNAs targeting the indicated genes or a scrambled control. Forty-eight hours after transfection, mScarlett-positive single cells were isolated via fluorescence-activated cell sorting (FACS) into RPMI-1640 medium supplemented with 20% FBS. Single clones were validated for gene deletion by immunoblotting using anti-UFM1, anti-UFSP2 or anti-DLAT antibodies before being pooled for downstream analysis.

To generate stable cell lines overexpressing UFM1, UFSP2 or DLAT variants, cDNAs of interest were integrated into the Rogi2 safe-harbor locus^62^. Cells were subjected to selection with 1 µg/mL puromycin (Thermo Fisher Scientific, NC9138068) starting 48 hours after transfection until the parental non-transfected population was eliminated. Protein overexpression was confirmed by immunoblotting using anti-UFSP2, anti-DLAT, or anti-Flag antibodies. All DNA oligonucleotides were synthesized by IDT.

To transiently overexpress UBA5, UFC1, UFL1, DDRGK1, PDHA1, PDHB, DLAT and DLD, their cDNAs, acquired from McDermott Center Sequencing Core at UT Southwestern, were cloned into pCDNA5 vector (kindly provided by Dr. Gerta Hoxhaj).

The specific gRNA sequences used were as follows:

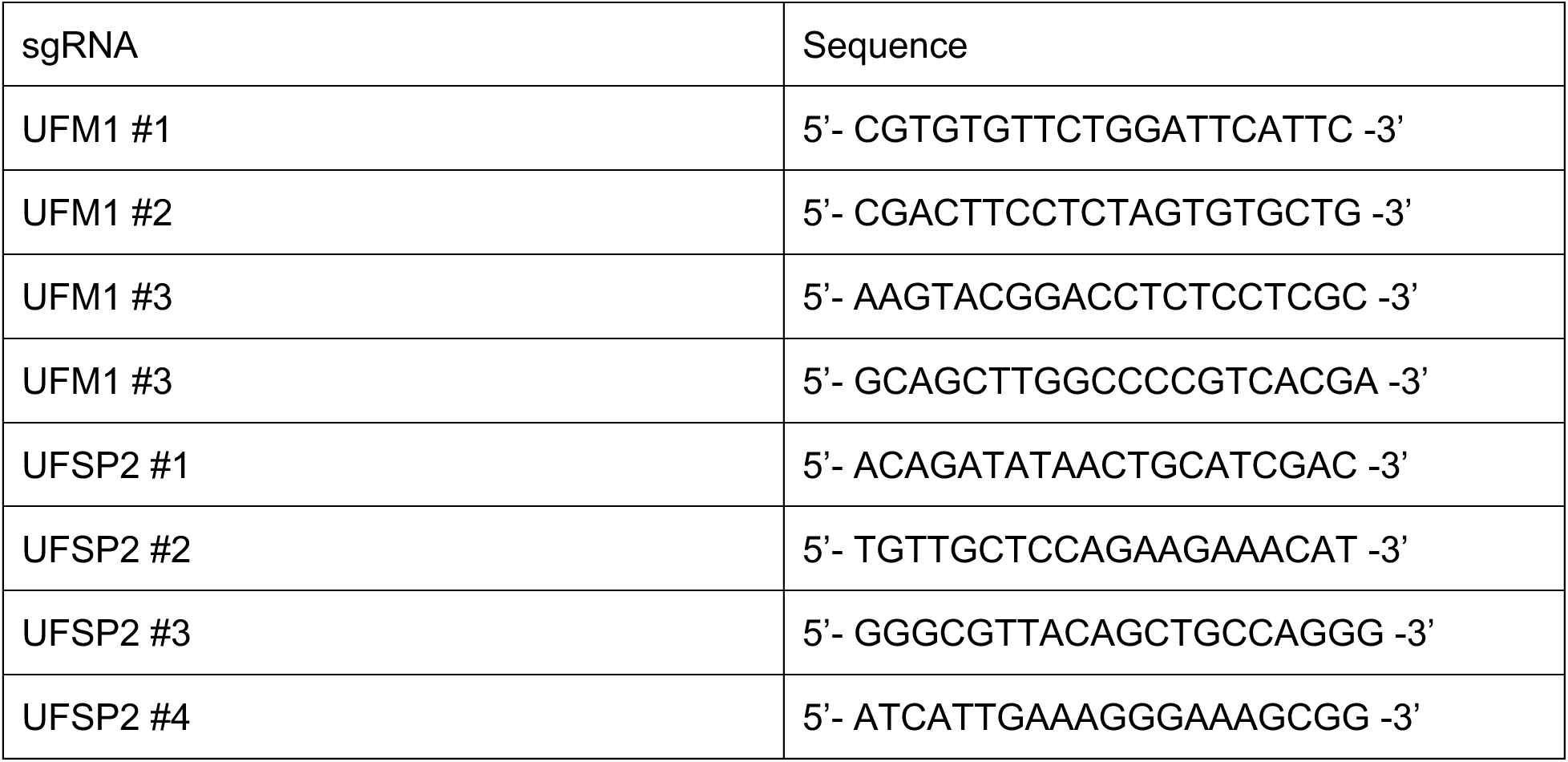
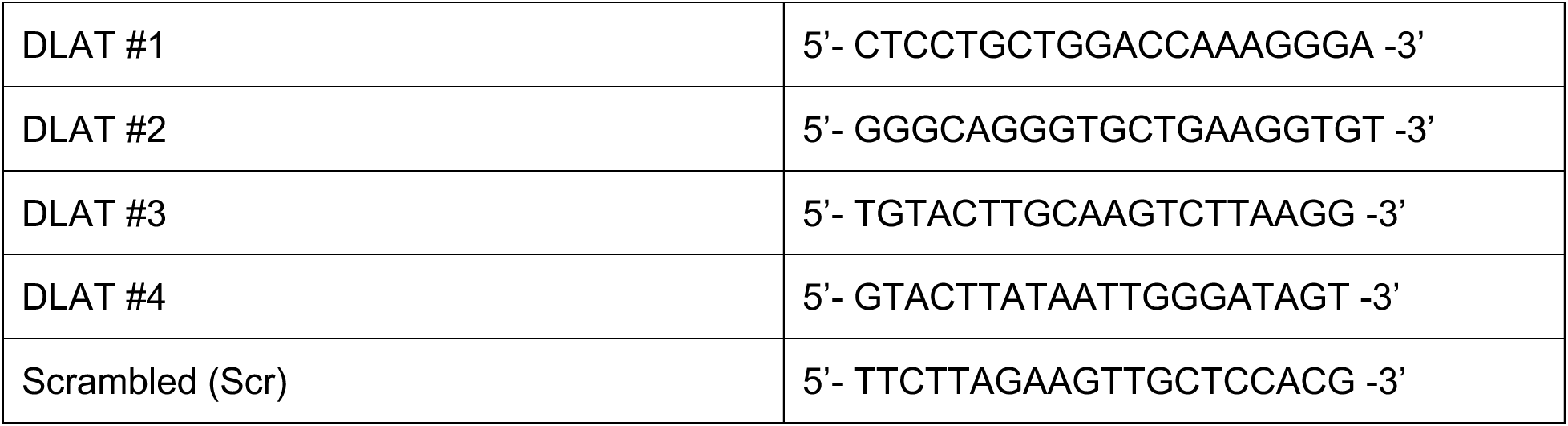

### Immunoblotting

RIPA buffer (Boston BioProducts, BP-115) containing proteinase and phosphatase inhibitors (Thermo Fisher Scientific, 78444) was used to lyse cells. Samples were centrifuged at 20,160 x g at 4°C for 10 minutes followed by protein quantification using the DC Protein Assay Kit. Equal amounts of protein were loaded on the gel, followed by transfer to PVDF membranes (Thermo Fisher Scientific, 88518). Membranes were blocked with 5% BSA in Tris-HCl buffer (TBS) containing 0.1% Tween-20 (TBST), subsequently incubated with primary antibodies diluted in filtered TBST containing 5% BSA overnight in the cold room. The membranes were washed with TBST 3 times for 5 minutes at room temperature followed by incubation with the secondary antibody (Cell Signaling Technology, 7074, 7076) diluted in 5% non-fat milk in TBST for 1 hour at room temperature. Membranes were washed three times for 5-10 minutes with TBST at room temperature before being incubated with Pierce ECL (Thermo Fisher Scientific, PI32106) for 2 minutes. Autoradiography films were used to detect the signals.

### Immunoprecipitation-coupled Mass spectrometry (IP-MS)

### Immunoprecipitation

To profile UFMylated proteins (the “UFMylome”) accumulating in UFSP2-depleted cells, we engineered *△*UFSP2 HEK 293T cells stably expressing either Flag-tagged wild-type UFM1 (UFM1^WT^) or a non-conjugatable Flag-tagged mutant (UFM1^ΔGSC^) as a negative control. The *△*GSC mutation removes the C-terminal “GSC” tripeptide, including the glycine residue essential for UFM1 conjugation. To enhance the UFMylation of targeted proteins, UFM1-system enzymes- including UBA5, UFC1, UFL1 and DDRGK1 were transiently overexpressed as previously described^63^. Both free and conjugated UFM1 species were immunoprecipitated using Flag-M2 resin and subsequently analyzed by mass spectrometry to identify candidate UFMylated proteins. Pulldown samples were maintained at -80 °C prior to analysis.

### Sample preparation

On the day of analysis, samples were removed from -80°C, then 1,220 μL LC-MS grade methanol was added to each tube and vortexed vigorously for 10 seconds to precipitate protein. Samples were centrifuged at 14,000 g for 2 minutes at 5°C to pellet precipitated protein. The resulting supernatant was then decanted to waste, ensuring not to disturb the protein pellet. Protein pellets were resuspended in 50 μL of 8M urea in 100 mM Tris with 10 mM TCEP and 40 mM chloroacetamide, then vortexed at ambient temperature for at least 20 minutes to resolubilize protein. 1 μL of 1 mg/mL LysC prepared per manufacturer’s instruction (VWR, Radnor, PA) was added to each sample, then allowed to incubate at ambient temperature for four hours while gently rocking. After incubation, 150 μL 100 mM Tris HCl pH 8.0 was added to each sample, gently pipetting up and down to mix. 2.5 μL 0.4 mg/mL trypsin (Promega, Madison, WI) was added to each sample, then samples were allowed to digest overnight at ambient temperature while rocking. The following day, the digestion reaction was terminated by adding 40 μL 10% TFA in H2O. Samples were then centrifuged at 14,000 g for 2 minutes at ambient temperature to pellet insoluble material. Resulting supernatants were desalted using Strata-X 33 μm polymeric reversed phase SPE cartridges (Phenomonex, Torrance, CA). Desalted peptides were dried down in a vacuum centrifuge (Thermo Fisher Scientific, Waltham, MA). Peptides were resuspended in 0.2% formic acid in water to determine peptide concentrations via NanoDrop One Microvolume UV-Vis spectrophotometer (Thermo Fisher Scientific, Waltham, MA), then vialed for instrument analysis.

### Proteomics Analysis

Samples were analyzed using a Dionex UltiMate 3000 RSCLnano liquid chromatography system (Dionex, Sunnyvale, CA) coupled to an Orbitrap Eclipse Tribrid mass spectrometer (Thermo Fisher Scientific, San Jose, CA). Mobile phase A consisted of 0.2% formic acid in H_2_O; mobile phase B consisted of 0.2% formic acid in an 80:20 ACN:H_2_O solution. Gradient elution was performed at a flowrate of 0.350 μL/minute. 500 ng of peptides were loaded onto a 75 μm i.d. column with 1.7 μm, 130 Å pore sized, Bridged Ethylene Hybrid (BEH) C18 particles (Waters, Milford, MA), packed in-house to a length of 30 cm and heated to 50 °C during analysis. For mass spectrometer analysis, in positive mode, MS1 scans were acquired from 0 to 60 minutes every 1.5 seconds with a scan range of 300-1350 m/z at a resolution of 60,000 in the Orbitrap; normalized AGC target was set to 200% (8e5 ions), maximum injection time was 50 ms, and RF lens (%) was set to 30. Precursor ions were isolated from a 0.9 Da window in the quadrupole; data-dependent HCD MS/MS scans were performed with 28% normalized collision energy, normalized AGC target of 200% (equivalent to 1e5 ions), and were collected in the Orbitrap from a starting first mass of 150 m/z, with a maximum injection time of 25 ms and dynamic exclusion period set to 10 seconds.

### Data Analysis

Raw proteomic data files were processed by MaxQuant 1.5.2.81^64^ and searched against the UniProt human database downloaded June 2022 including isoforms. Default MaxQuant parameters were used to process, along with the following parameters: label-free quantification (LFQ) calculated with a minimum ratio of 1; match between runs enabled; MS/MS not required for LFQ comparisons; and VG of lysine (+156.0898 Da) as variable modification. For data processing: from the generated MaxQuant output, protein identifications were removed that were indicated to be identified by site only, corresponded to reverse sequences, and/or to be potential contaminants by MaxQuant. Protein identifications that generated an intensity value of zero in 50% or more of the analyzed samples were also removed. Missing quantitative values among the remaining protein groups were imputed, log_2_-transformed, and statistically analyzed using Argonaut2^65^.

### Data Availability

The raw data has been deposited to the MassIVE database under the accession number MSV000100954. The files can be accessed through doi:10.25345/C5NC5SS4V with the password “UFM1proteomics”.

### mtDNA copy number analysis

Total DNA was extracted from cells using DNeasy Blood and Tissue kit (Qiagen, 69504) according to the manufacturer’s instructions. DNA concentrations were adjusted to 10 ng/µL and subjected to quantitative PCR analysis. Primers targeting COX2 (mitochondrial DNA) and H4C3 (nuclear DNA) were used to quantify mtDNA and nuclear DNA, respectively. The Ct value of COX2 was normalized to the Ct value of H4C3 for each sample. Three technical replicates were performed for each biological sample, and the mean Ct value was used for the relative mtDNA copy number analysis using the 2^−ΔΔCt^ method. The mtDNA abundance of the control group was set to 100%.

COX2-Forward: CCGTCTGAACTATCCTGCCC

COX2-Reverse: GCCGTAGTCGGTGTACTCGT

H4C3-Forward: GGGATAACATCCAGGGCATT

H4C3-Reverse: CCCTGACGTTTTAGGGCATA

### Mitochondrial membrane potential analysis

Mitochondrial membrane potential was assessed as previously described^66^. In brief, cells were stained with 20 nM MitoTracker Deep Red (Invitrogen, M46753) for 30 minutes, then pelleted and resuspended in FACS buffer containing 1x phosphate buffered saline (PBS) (Ca^2+^/Mg^2+^ free), 2.5 mM EDTA, 2.5 mM HEPES, pH 7.0, 1% heat-activated Fetal Bovine Serum (FBS), 1% penicillin-streptamycin (Pen-Strep) prior to flow cytometry analysis. Data were analyzed using FlowJo software.

### Seahorse XFe96 Respirometry

Oxygen consumption rates were measured using an XFe96 Extracellular Flux Analyzer (Agilent Technologies). 15,000 cells were seeded in a 96-well Seahorse plate for 16-18 hours. Cells were then washed twice with DMEM containing 10 mM glucose, 1 mM pyruvate, 2 mM glutamine, 1% Pen-Strep and no bicarbonate followed by at least 30 minutes of incubation in a CO_2_-free incubator at 37°C before the analysis. Oxygen consumption rates were normalized to the cell number in each well.

### Blue-native polyacrylamide gel electrophoresis (BN-PAGE)

To analyze ETC complex assembly, we followed a published protocol^67^. Control, ΔUFM1 and ΔUFSP2 cells were lysed on ice in digitonin buffer containing PBS supplemented with 50 ug/ml digitonin, and 1x protease and phosphatase inhibitor (Thermo Fisher Scientific, 78444) for 10 minutes, and centrifuged at 10,000 x g for 10 min at 4°C. Pellets were resuspended in digitonin buffer, and crude mitochondria were isolated via centrifugation at 20,000 x g for 10 minutes at 4°C. 100 µg mitochondrial fractions were solubilized in 40 µL of sample buffer (Invitrogen, BN2003) supplemented with 2% digitonin for 30 minutes on ice. Following centrifugation (20,000 x g, 10 minutes, 4°C), 36 µL of the supernatant was mixed with 4 µL of 5% G-250 sample additive (Invitrogen, BN2004).

For BN PAGE, 10 µg of protein was loaded onto 3-12% Bis-Tris NativePAGE gels (Invitrogen, BN1001BOX). The gels were run at 150 V, 4°C for 30 minutes using 1x anode buffer (Invitrogen, BN2001) and 1xcathode buffer containing 1.25% cathode additive (Invitrogen, BN2002). The cathode buffer was then replaced with 1x anode buffer, and electrophoresis continued at 150 V, 4°C for 90 minutes. Prior to transfer, gels were equilibrated for 15 min in Twobin buffer containing 25 mM Tris, 192 mM glycine, 20% v/v methanol, 0.1% SDS. Proteins were transferred to membranes using a Trans-Blot Turbo system (Bio-Rad) at 25 V for 30 min.

Membranes were fixed in 8% acetic acid (5 min), rinsed with water, and sequentially washed with 100% methanol and water to remove residual G-250 dye. After blocking in 5% BSA, membranes were incubated with primary antibodies prepared in 5% BSA TBST against NDUFB8 (Abcam ab192878, 1:3,000), SDHA (CST 11998S, 1:1,000), UQCRC2 (Abcam ab14745, 1:1,000), MTCO1 (Abcam ab154477, 1:3,000), or ATP5A (Proteintech 66037-1-Ig, 1:5,000) overnight. HRP-conjugated anti-rabbit (Fisher Scientific AP182PMI, 1:10,000) and anti-mouse (Cell Signaling Technology, 7076P2) secondary antibodies.

### Tandem Mass Tagging Mass Spectrometry (TMT-MS)

Control, ΔUFM1 and ΔUFSP2 HeLa cells were rinsed twice with ice-cold PBS and lysed in a buffer containing 5% SDS in 50 mM TEAB, supplemented with protease and phosphatase inhibitors. After 15 min incubation, proteins were quantified using DC Protein Assay Kit (Bio-Rad, 5000111) and normalized by the same total input (50 µg). Following reduction and alkylation, proteins were digested overnight with trypsin on S-Trap column (Protifi). Eluted peptides were labeled with TMT10plex Isobaric Mass Tagging Kit (Thermo), pooled and desalted (Oasis HLB, Waters, and dried in a SpeedVac. The dried sample was reconstituted in 0.1% TFA buffer containing 2% acetonitrile and then diluted down to ∼1 ug of peptides before injection. Analysis was performed on a Thermo Orbitrap Eclipse MS coupled to an Ultimate 3000 RSLC-Nano liquid chromatography system^68^.

Mitochondrial proteins within each genotype were ranked based on their median abundance across replicates in the whole-cell lysate TMT-MS dataset. The distribution of abundance ranks for mitochondrial proteins was compared across genotypes using Kruskal-Wallis non-parametric tests.

### Stable isotope tracing

For [U-^13^C]glucose tracing, cells were cultured in glucose-free RPMI medium (Sigma, R1383-L) supplemented with 11 mM [U-^13^C]glucose (Cambridge Isotope Laboratories, CLM481-0.25) and 10% dialyzed FBS (Gemini Bio-Products, 100108). For glutamine tracing, cells were cultured in glutamine-free RPMI medium (Sigma, R0883) supplemented with 2 mM [U-^13^C]glutamine (Cambridge Isotope Laboratories, CLM-1822-0) and 10% dialyzed FBS. Specific tracing durations were indicated in the figure legends. Extracted metabolites were analyzed via Gas chromatography/mass spectrometry (GC/MS) as described below.

### Gas chromatography/mass spectrometry (GC/MS)

To extract metabolites, cells were rinsed with ice-cold saline, harvested by scraping, and subjected to three freeze-thaw cycles. Following 1 min of vortexing, lysates were centrifuged at 20,160 x g at 4°C for 15 min. Supernatants were transferred to clear microcentrifuge tubes and dried overnight in a SpeedVac concentrator. The dried metabolites were resuspended in 30 µL of anhydrous pyridine containing 10 mg/mL methoxyamine. After brief vortexing and centrifugation, supernatants were transferred to GC/MS autoinjector vials and incubated at 75°C for 15 min. Derivatization was initiated by adding 70 µL N-(tert-butyldimethylsilyl)-N-methyltrifluoroacetamide (MTBSTFA), followed by a1-hour incubation at 75°C. A 1 µL of each sample was injected into an Agilent 5973N or 5975C Mass Spectrometer coupled to Agilent 6890 or 7890 Gas Chromatograph. Data analysis was performed using El-MAVEN, and natural abundance was corrected using MATLAB as previously described^69^.

### Acetyl-CoA measurement

For [U-^13^C] glucose tracing, cells (90% confluent, 10-cm dish) were incubated for 30, 60, and 180 min in DMEM medium (Sigma, D3060) supplemented with 4 mM glutamine and 11 mM [U-^13^C]glucose. For [U-^13^C] pyruvate tracing, cells (90% confluent, 10-cm dish) were incubated for 30 min in DMEM medium (Sigma, D3060) supplemented with 4 mM glutamine and 1 mM [U-^13^C]sodium pyruvate. Following incubation, cells were washed with PBS and lysed in 1 mL of ice-cold 10% trichloroacetic acid (TCA). Lysates were sonicated (12 × 0.5-s pulses) and clarified by centrifugation at 20,000g, 10 min, 4 °C. The resulting supernatants were loaded onto Oasis solid-phase extraction (SPE) columns (Waters, 186000679). After a 5-min incubation, the samples were passed through the columns using a vacuum manifold. Columns were washed with 1 ml of water, and acetyl-CoA was eluted with 300 ul of 25 mM ammonium acetate in methanol.

LC-MS/MS analysis was performed using an AB SCIEX QTRAP 5500 liquid chromatograph/triple quadrupole mass spectrometer as previously described^70^. Chromatographic separation was achieved on a Nexera Ultra-High-Performance Liquid Chromatograph (UHPLC) system (Shimadzu Corporation) using a SeQuant^®^ ZIC^®^-pHILIC HPLC column (150 × 2.1 mm, 5 µm, polymeric). The mass spectrometer utilized an electrospray ionization (ESI) source in multiple reaction monitoring (MRM) mode.

Mobile phase A consisted of 10 mM ammonium acetate in water (pH 9.8, adjusted with ammonium hydroxide) and mobile phase B consisted of 100% acetonitrile (ACN). The gradient elution was: 0–15 min, linear gradient 90–30% B, then the column was washed with 30% B for 3 min before reconditioning it for 6 min using 90% B. The flow rate was 0.25 mL/min, and the injection volume was 20 µL. MRM data were analyzed using Analyst 1.6.3 software (SCIEX). The MRMs used for acetyl-CoA isotopomers were: M+0 (Q1/Q3: 810/303, CE: 42), M+1 (Q1/Q3: 811/303 and 811/304, CE: 42), and M+2 (Q1/Q3: 812/303, 812/304, and 812/305, CE: 42).

### Pyruvate dehydrogenase (PDH) activity assays

PDH enzymatic activity was measured using a microplate assay kit (Abcam, ab109902) according to the manufacturer’s instructions. Briefly, cells (90% confluent, 15-cm plate) were rinsed with PBS twice, harvested by scraping, and resuspended in 9 volumes of PBS. To normalize protein loading, a 45 µL aliquot of the cell suspension was lysed in 5 µL lysis buffer containing 2% SDS. Following a 10 min incubation at room temperature (RT) and centrifugation at 20,000 x g, 10 min, protein concentration was determined using a DC protein assays (Biorad).

Samples were then normalized by protein concentration, treated with 1/10 volume of detergent, and incubated on ice for 10 min to solubilize membranes. The lysate was clarified by centrifugation at 1,000 x g for 10 min at 4°C. The resulting supernatant (200 µL per well) was loaded onto the assay plate and incubated for 3 hours at RT. Wells were washed twice with 300 µL 1x stabilizer buffer was used to wash the wells twice before adding 1x reaction mixture. PDH activity was monitored kinetically every 30 seconds for 60 min (120 cycles).

### Validation of UFMylated candidates

Flag-Strep-tagged wild-type PDHA1, PDHB, DLAT, and DLD were transiently expressed in ΔUFM1/ΔUFSP2 293F cells stably expressing either UFM1^WT^ or UFM1^ΔGSC^. Transfections were performed using polyethylenimide 25,000 (PEI) at a DNA:PEI ratio of 1:3. Approximately 100×10^6^ 293F cells were cultured in FreeStyle^TM^ 293 medium at 37°C, 8% CO_2_ for 48 hours, then harvested by centrifugation (1,000 x g for 20 min, 4°C).

Cell pellets were lysed in a buffer containing 20 mM Tris-HCl pH 7.5, 150 mM NaCl, 1 mM EDTA, 1% NP-40, and protease inhibitors (1 µg/ml leupetin, 1 µg/ml pepstatin, 1 mM benazamidine HCl). Lysates were subjected to three freeze-thaw cycles using liquid nitrogen and a 37°C water bath, followed by centrifugation at 18,000 x g for 15 min at 4°C. Protein concentrations in the resulting supernatants were determined using a DC protein assay kit.

For input control, normalized 50 µL of 2 µg/µL aliquots of the supernatants were mixed with 10 µL 6x Laemmli buffer and boiled at 95°C for 5 min. For affinity purification, 10 mg of total protein was incubated with 50 µL 50% Strep-Tactin XT Sepharose chromatography resin (Sigma, GE29401324) on a rotator for 2 hours at 4°C. The resin was collected by centrifugation at 1,000 x g for 30 seconds at 4°C. After aspirating the supernatant, the resins were washed five times with 1 ml TBST followed by centrifugation at 1,000 x g at 4°C for 30 seconds. Bound proteins were eluted by three serial additions of 25 µL 2x Laemmli buffer followed by incubation at 95°C for 5 min. Samples were subsequently analyzed by immunoblotting or processed for mass spectrometry to identifiy KGV-containing peptides as described below.

### KGV-containing peptide identification

Samples were digested overnight with trypsin (Pierce) following reduction and alkylation with DTT and iodoacetamide (Sigma–Aldrich). Following solid-phase extraction cleanup with an Oasis HLB µelution plate (Waters), the resulting peptides were reconstituted in 10 uL of 2% (v/v) acetonitrile (ACN) and 0.1% trifluoroacetic acid in water. 5 uL of the peptides were injected into an Orbitrap Fusion Lumos mass spectrometer (Thermo) coupled to an Ultimate 3000 RSLC-Nano liquid chromatography systems (Thermo). Samples were injected into a 75 μm i.d., 75-cm long EasySpray column (Thermo), and eluted with a gradient from 0-28% buffer B over 90 min. Buffer A contained 2% (v/v) ACN and 0.1% formic acid in water, and buffer B contained 80% (v/v) ACN, 10% (v/v) trifluoroethanol, and 0.1% formic acid in water. The mass spectrometer was operated in positive ion mode with a source voltage of 2.5 kV and an ion transfer tube temperature of 300 °C. MS scans were acquired at 120,000 resolution in the Orbitrap and up to 10 MS/MS spectra were obtained in the Orbitrap for each full spectrum acquired using higher-energy collisional dissociation (HCD) for ions with charges 2-7. Dynamic exclusion was set for 25 s after an ion was selected for fragmentation.

Raw MS data files were analyzed using Proteome Discoverer v3.0 (Thermo), with peptide identification performed using Sequest HT searching against the human reviewed protein database from UniProt (downloaded January 4, 2024, 20354 entries). Fragment and precursor tolerances of 10 ppm and 0.6 Da were specified, and three missed cleavages were allowed. Carbamidomethylation of Cys was set as a fixed modification, with oxidation of Met and Val-Gly modification of Lys set as variable modifications. The false-discovery rate (FDR) cutoff was 1% for all peptides. Peptide peak intensities were used as peptide abundance values.

### Immunoprecipitation

Flag-Strep-tagged wild-type DLAT (WT) and its lysine-to-arginine variants (K118R, K363R, and K547R) were stably expressed in control, ΔUFM1 and ΔUFSP2 HeLa cells as described above. Cells (90% confluent, 5 15-cm plates) were harvested by scraping in ice-cold PBS and pelleted at 4°C. The pellets were resuspended in lysis buffer containing 20 mM Tris-HCl pH 7.5, 150 mM NaCl, 1 mM EDTA, 1% NP-40, and protease inhibitors (1 µg/ml leupetin, 1 µg.ml pepstatin, 1 mM benazamidine HCl). Lysates were subject to three freeze-thaw cycles between liquid nitrogen and a 37°C water bath, followed by centrifugation at 18,000 x g for 15 min at 4°C.

Supernatants were collected and protein concentrations were determined using a DC protein assay kit. For input control, normalized 50 µL aliquots of the supernatants were mixed with 10 µL 6x Laemmli buffer and boiled at 95°C for 5 min. For affinity purification, approximately 10 mg protein from the supernatant was incubated with 50 µL 50% Strep-Tactin XT Sepharose chromatography resin (Sigma, GE29401324) on a rotator at 4°C for 2 hours. The samples were centrifuged at 1,000 x g at 4°C for 30 seconds. The supernatants were aspirated and the pellets were washed with 1 mL TBST followed by centrifugation at 1,000 x g at 4°C for 30 seconds. After five washes, 25 µL 2x Laemmli buffer was added followed by 5 min of incubation at 95°C to elute the samples three times. Western blots were performed as described in **Immunoblotting.**

### Statistical analysis

Figures were prepared and statistics were calculated using GraphPad PRISM and R. Statistical calculation details are indicated in the figure legends for each figure. NS, not significant (P > 0.05).

